# Species-specific deployment of Runx2 isoforms and differential regulation of target genes during avian jaw development and evolution

**DOI:** 10.1101/2021.05.25.444052

**Authors:** Spenser S. Smith, Daniel B. Chu, Tiange Qu, Tiffany Huang, Austen J. Lucena, Goutam Krish, Richard A. Schneider

## Abstract

Developmental regulation of bone formation in the jaw skeleton is essential to species-specific adaptation. The jaws are derived from neural crest mesenchyme (NCM), a progenitor population that directs skeletal patterning by exerting temporal and spatial control over molecular and cellular programs for osteogenesis. One important NCM-mediated gene is *Runx2*, which is a transcription factor required for osteoblast differentiation. RUNX2 protein binds many target genes involved in the deposition and resorption of bone. To determine the extent to which changes in *Runx2* structure, function, and expression underlie the evolution of the jaw skeleton, we compare *Runx2* across vertebrates and within birds. *Runx2* contains two alternative promoters, tandem repeats of glutamine and alanine with variable lengths in different species, a conserved DNA-binding domain, an exon that is alternatively spliced, as well as two possible C-termini. Such alternative splicing produces eight potential isoforms that show distinct stage- and species-specific patterns in the jaw primordia of chick, quail and duck embryos. We also find that certain isoforms are strongly induced by TGFβ signaling whereas others are not. Overexpressing *Runx2* isoforms in NCM reveals that some are transcriptionally activating, while others are repressive. But context appears to be relevant since species-specific polymorphisms in the promoter of target genes like *Mmp13*, can modulate the effects of different isoforms. Overall, our study indicates that the structure and species-specific deployment of *Runx2* isoforms affect the transcriptional activity of target genes in ways that may have played a generative and regulatory role in the evolution of the avian jaw skeleton.

## INTRODUCTION

The craniofacial skeleton is one of the most highly adapted structures of vertebrates. Its anatomy often evolves in ways that seem perfectly suited to meet functional and ecological demands. To understand how the craniofacial skeleton becomes modified during evolution, we have been assaying for species-specific differences in molecular and cellular programs underlying skeletogenesis. Experimental work spanning almost a century has shown that neural crest mesenchyme (NCM), which is the precursor population that gives rise to the cartilages and bones of the jaws and face, is the primary source of species-specific pattern (Harrison, 1935; de Beer, 1947; Hörstadius, 1950; Harrison, 1969; Hall and Hörstadius, 1988; Noden and Schneider, 2006; Fish and Schneider, 2014b; Schneider, 2018b; Woronowicz and Schneider, 2019). For example, embryonic transplants of presumptive NCM from quail or chick into duck produce short quail- and chick-like beaks and jaw skeletons on duck hosts, and long duck-like bills on quail hosts (Schneider and Helms, 2003; Tucker and Lumsden, 2004; Schneider, 2005; Eames and Schneider, 2008; Solem et al., 2011; Woronowicz et al., 2018). To accomplish such a complex and profound task, NCM exerts spatial and temporal control over the expression of genes that mediate the induction, deposition, and resorption of bone (Merrill et al., 2008; Hall et al., 2014; Ealba et al., 2015; Schneider, 2015). One gene in particular, *Runt related transcription factor 2* (*Runx2*), shows intriguing species-specific patterns with quail expressing higher endogenous levels than duck, coincident with their smaller jaw size. *Runx2* is considered a master regulator of osteogenesis as evidenced by its effects on osteoblasts and the loss of bone in *Runx2* mutant mice (Ducy et al., 1997; Komori et al., 1997; Otto et al., 1997; Karsenty et al., 1999; Ducy, 2000; Teplyuk et al., 2008; Komori, 2010b; Adhami et al., 2015; Vimalraj et al., 2015; Takarada et al., 2016; Shirai et al., 2019). Over-expressing *Runx2* prematurely or at higher levels can reduce the size of the jaw skeleton (Eames et al., 2004; Hall et al., 2014). Exactly how higher levels of *Runx2* expression relate to smaller jaw size is unclear, but presumably mechanisms involve the differential activation and/or repression of target genes.

RUNX2 is an essential effector of the *Transforming Growth Factor-Beta* (*Tgfβ*) superfamily signaling pathway (Lee et al., 2000; Alliston et al., 2001; Lee et al., 2002; Ito and Miyazono, 2003; Kang et al., 2005; Derynck et al., 2008) and directly binds a broad array of target genes involved in the deposition or resorption of bone including *Osteocalcin* (*Ocn*), *Collagen type 1* (*Col1a1*), as well *Matrix metalloproteinases* such as *Mmp2* and *Mmp13* (Jimenez et al., 1999; Xiao et al., 1999; Kern et al., 2001; Zaidi et al., 2001; Eames et al., 2003; Otto et al., 2003; Schroeder et al., 2004; Wang et al., 2004; Pratap et al., 2005; Schroeder et al., 2005; Makita et al., 2008; Komori, 2010a). In a generalized form, *Runx2* contains a distal promoter known as P1 (or MASNS after its initial amino acid sequence) and a proximal promoter known as P2 (or MRIVP), a conserved DNA binding Runt Homology Domain (RHD), tandem repeats of glutamine (Q) and alanine (A), as well as two C-termini and other exons that are alternatively spliced to generate multiple isoforms (**Figure 1A**) (Coffman, 2003; Rennert et al., 2003; Levanon and Groner, 2004; Terry et al., 2004; van Wijnen et al., 2004; Stock and Otto, 2005; Vimalraj et al., 2015; Mevel et al., 2019; Kim et al., 2020). Changes to *Runx2* gene structure including the ratio of poly Q to poly A, alternative splicing of isoforms, and splice-site mutations associate with normal variation as well as abnormal defects and disease in the skeleton (Sohn et al., 1994; Harada et al., 1999; Xiao et al., 1999; Banerjee et al., 2001; Chen et al., 2002; Choi et al., 2002; Yoon et al., 2002; Albrecht et al., 2004; Vaughan et al., 2004; Kim et al., 2006; Achari et al., 2008; Makita et al., 2008; Pasche et al., 2010; Sun et al., 2011; Morrison et al., 2013; Okura et al., 2014; Mastushita et al., 2015; Jaruga et al., 2016; Shibata et al., 2016; Knobloch et al., 2019). Yet despite a well-established role for *Runx2* as a key transcriptional regulator of osteogenesis, the extent to which species-specific changes in its structure, function, and/or expression underlie the evolution of the jaw skeleton remains poorly understood.

**Figure 1.**
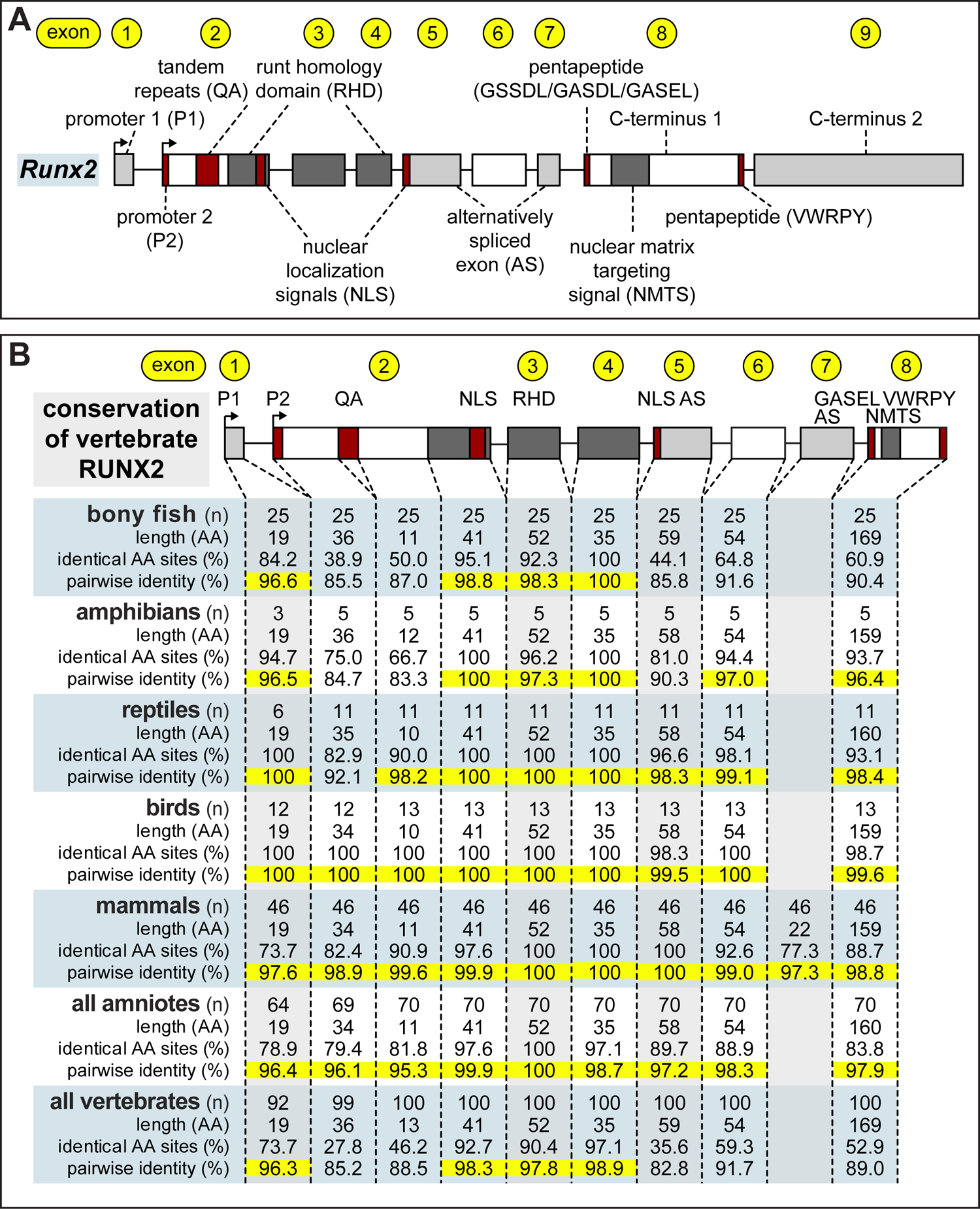
Structure and conservation of RUNX2 domains across vertebrates. **A)** In a generalized form, the *Runx2* gene contains two alternative promoters and up to nine exons that encode for distinct functional domains as shown schematically. RUNX2 protein domains can include tandem repeats of glutamine (Q) and alanine (A), which vary in repeat length depending on the taxon; a runt homology domain (RHD), which gives the gene its name and is responsible for DNA-binding and protein-protein interactions; nuclear localization signals (NLS); alternatively spliced (AS) exons; pentapeptide domains; a nuclear matrix targeting signal (NMTS); and two alternative C-termini. **B)** For the most part, RUNX2 functional domains are highly conserved across vertebrates as shown in a similarity matrix of amino acid (AA) sequences. However, the QA domain and C-terminus 2 are highly variable (and are not included in the similarity matrix). For purposes of analysis, exon two was divided into three components: the region upstream of the QA repeat, the region downstream of the QA repeat to the RHD, and the region containing part of the RHD. Length refers to the number of AA within a given domain. Identical AA sites are defined as the number of AA that are identical at each AA position within a region. Pairwise identity is calculated by making a pairwise comparison between each AA at each position and dividing the number of identical pairwise AA by the number of total pairwise comparisons. Pairwise identities over 95% are highlighted in yellow.

To address this issue, we first examine the architecture of *Runx2* at a high level across vertebrates and map the major transitions in *Runx2* gene evolution. Our goal is to distinguish between domains that are variable or conserved. Then we perform a series of *in vitro* and *in vivo* functional analyses of *Runx2* primarily using a three-taxon comparison in which two taxa are more closely related than either are to a third, since this is considered a robust strategy for making parsimonious inferences about evolution (Nelson and Platnick, 1991; Mavrodiev et al., 2019; Rineau et al., 2020). We exploit duck, chick, and quail in particular, because their jaw skeletons differ in size (from relatively large to small), their genomes are sequenced and annotated, and their embryos can be easily matched using the Hamburger and Hamilton (HH) staging system (Hamburger and Hamilton, 1951; Koecke, 1958; Padgett and Ivey, 1960; Zacchei, 1961; Hamilton, 1965; Yamashita and Sohal, 1987; Le Douarin et al., 1996; Ricklefs and Starck, 1998; Starck and Ricklefs, 1998; Nakane and Tsudzuki, 1999; Schneider and Helms, 2003; Stern, 2005; Lwigale and Schneider, 2008; Sauka-Spengler and Barembaum, 2008; Jheon and Schneider, 2009; Ainsworth et al., 2010; Mitgutsch et al., 2011; Fish and Schneider, 2014a; Gammill et al., 2019; Chu et al., 2020; Smith et al., 2020). As Galliformes, chick and quail are separated by approximately 50 million years, and they diverged from a common ancestor with Anseriformes, which include duck, around 100 million years ago (Pereira and Baker, 2006; Hackett et al., 2008; Kan et al., 2010).

Our comparative analyses reveal that RUNX2 is extremely conserved across vertebrates with the exception of the QA repeats and second C-terminus. Such conservation suggests that instead of major modifications within individual domains, spatiotemporal changes to the levels of *Runx2* expression and/or alternative splicing, could serve as a primary mechanism for skeletal evolution by differentially regulating target genes. In support of this hypothesis, our spatial and temporal analyses of *Runx2* gene and protein expression in the developing jaw primordia of chick, quail, and duck uncover species-specific differences at embryonic stages when the bony skeleton is undergoing deposition and resorption. Moreover, by sequencing *Runx2* mRNA transcripts present in the jaw primordia during these stages, we detect the presence of eight *Runx2* isoforms. A subsequent RNA isoform sequencing (Iso-Seq) experiment reveals species-specific differences in the proportions and spatial localization of these isoforms, which we then show can vary over time and become differentially expressed in response to recombinant (r) TGFβ1. To test the ability of these isoforms to regulate target genes, we perform overexpression experiments *in vitro* and *in ovo* and find that some isoforms are activating while others are repressive, and their effects vary among species. As a proof-of-principle, we focus on the regulation of the *Mmp13* promoter by RUNX2. We test the functional consequences of two single nucleotide polymorphisms (SNPs) directly downstream of the RUNX2 binding element within the *Mmp13* promoter that distinguish quail and chick from duck. Switching these SNPs between species alters *Mmp13* promoter activity presumably by affecting interactions with RUNX2 isoforms. Overall, our studies demonstrate that alterative splicing of *Runx2* is a key feature of avian jaw development and that the combinatorial, spatial, temporal, and/or species-specific expression of *Runx2* isoforms by NCM differentially regulates downstream target genes involved in the deposition and resorption of bone, and likely affects jaw anatomy during evolution.

## MATERIALS AND METHODS

### Genomic Analysis and Sequencing of Runx2

The National Center for Biotechnology Information (NCBI) GenBank database was queried using “runx2” as a search term resulting in 215 sequences from species with an annotated *Runx2* gene. Each sequence was aligned to human, mouse, and chick *Runx2* sequences as well as *Runx2* motif sequences to determine if the full length *Runx2* sequence was present. Incomplete sequences were excluded from further analysis. FastTree (Price et al., 2010) was used in Geneious (Version 2020.2.4, Geneious Prime, Auckland, New Zealand) to generate an unrooted tree of all of the full length RUNX2 amino acid (AA) sequences via the BLOSUM62 cost matrix (Henikoff and Henikoff, 1992).

To identify sequences of the *Runx2* promoters used by chick, quail, and duck, 5’ rapid amplification of cDNA ends (RACE) reactions were carried out using template switching oligos (Matz, 2003; Picelli et al., 2014; Bagnoli et al., 2018). In short, reverse transcription was performed using the Maxima H Minus First Strand cDNA Synthesis Kit (K1651, Thermo Fisher Scientific, Waltham, MA, USA) following the manufacturer’s directions with 2 μg of total RNA, 100 pmol of *Runx2* primer, and 50 pmol of template switching oligo. The cDNA synthesis reaction was carried out at 50°C for 30 min, 55°C for 20 min, cycled five times for 2 min at 55°C and 2 min at 42°C, cycled five times for 2 min at 60°C and 2 min at 42°C, cycled 10 times for 65°C for 2 min and 2 min at 42°C, and 85°C for 5 min. Amplification of cDNA ends was carried out using step-out PCR as previously described (Matz et al., 2003) with the Q5 Hot Start High-Fidelity DNA Polymerase (M0493L, NEB, Ipswich, MA).

To sequence *Runx2* isoforms, full length cDNA synthesis from RNA was carried out using Maxima H-reverse transcriptase following the manufacturer’s directions with 2 μg of total RNA and 100 pmol of d(T)_20_ VN primer. The cDNA synthesis reaction was conducted at 50°C for 30 min, 55°C for 10 min, 60°C for 10 min, 65°C for 10 min, and 85°C for 5 min. To amplify each isoform, primers were designed targeting promoter 1 (P1) and promoter 2 (P2) sequences identified from the 5’ RACE and C-terminus 1 (C1) and C-terminus 2 (C2). All primers are listed in **Supplemental Table 1**. Amplification was carried out using Q5 Hot Start High-Fidelity DNA Polymerase following the manufacturer’s directions adding 1 μl of cDNA per 100 μl cycling 30 times at 98°C for 10 sec, 65°C for 20 sec, and 72°C for 2 min. Amplicons were purified using GeneJET PCR Purification Kit (K0702, Thermo Fisher Scientific, Waltham, MA, USA) following the manufacturer’s directions and cloned using CloneJET PCR Cloning Kit (K1231, Thermo Fisher Scientific, Waltham, MA, USA) following the manufacturer’s directions. At least 10 colonies were sequenced for each primer pair.

### The Use of Avian Embryos

For all experiments, we adhered to accepted practices for the humane treatment of avian embryos as described in S3.4.4 of the *AVMA Guidelines for the Euthanasia of Animals: 2013 Edition* (Leary et al., 2013). Fertilized eggs of chicken (*Gallus gallus*), Japanese quail (*Coturnix coturnix japonica*), and white Pekin duck (*Anas platyrhynchos domestica*) were purchased from AA Lab Eggs (Westminster, CA) and incubated at 37.5°C in a humidified chamber (GQF Hova-Bator 1588, Savannah, GA) until they reached embryonic stages appropriate for analyses. Embryos were matched at equivalent stages the using the Hamburger and Hamilton (HH) staging system, a well-established standard that utilizes an approach based on external morphological characters, that is independent of body size and incubation time, and that can be adapted to other avian species such as quail and duck (Hamburger and Hamilton, 1951; Hamilton, 1965; Ricklefs and Starck, 1998; Starck and Ricklefs, 1998; Schneider and Helms, 2003; Lwigale and Schneider, 2008; Jheon and Schneider, 2009; Ainsworth et al., 2010; Mitgutsch et al., 2011; Fish and Schneider, 2014a; Smith et al., 2015).

### Histological Staining and Immunohistochemistry

Quail and duck embryos were collected at HH37 in 4% paraformaldehyde (PFA) (15714, Electron Microscopy Sciences, Hatfield, PA, USA) overnight at 4°C (Schneider, 1999; Schneider et al., 2001). To detect bone deposition in sections, embryos were dehydrated in methanol, embedded in paraffin, and cut into 10 µm sagittal sections. Sections were deparaffinized, rehydrated, and adjacent sections were stained with Milligan’s trichrome at room temperature as previous described (Presnell and Schreibman, 1997; Schneider et al., 2001; Eames and Schneider, 2005; Tokita and Schneider, 2009; Solem et al., 2011; Hall et al., 2014). Immunohistochemistry (IHC) was performed on adjacent sections. For antigen retrieval, sections were heated in a microwave to 95°C in 10 mM sodium citrate buffer for 10 min and were blocked for endogenous peroxidase activity with 3% hydrogen peroxide for 15 minutes. Sections were incubated with 1:500 RUNX2 rabbit polyclonal primary antibody (AB23981, Abcam, Cambridge, UK) overnight at 4°C. Sections were stained with 1:500 goat anti-rabbit Alexa Fluor 647 secondary antibody (A32733, Thermo Fisher Scientific, Waltham, MA, USA) was overnight at 4°C. 10 mg/ml Hoechst 33342 dye (62249, Thermo Fisher Scientific, Waltham, MA, USA) was used to stain nuclei. Lower jaw sections were imaged using a Nikon AZ100 C2 macroconfocal microscope (Nikon Instrument, Inc., Melville, NY) for IHC, and a Leica DM 2500 (Leica Microsystems, Inc. Buffalo Grove, IL) with a color digital camera system (SPOT Insight 4 Megapixel CCD, Diagnostic Instruments, Inc., Sterling Heights, MI) for Trichrome.

### RNA Extractions and Quantitative PCR

Lower jaws were dissected from chick, quail, and duck embryos and total RNA was extracted using the RNeasy Plus Mini Kit (74136, Qiagen, Hilden, Germany) following the manufacturer’s protocol. Lower jaws were resuspended in 600 μl of RTL plus buffer supplemented with 1% β-mercaptoethanol (M3148-100ML, MilliporeSigma, Burlington, MA, USA) and Reagent DX (19088, Qiagen, Hilden, Germany). HH31 and HH34 lower jaws were processed in a Bead Mill 24 Homogenizer (15-340-163, Fisher Scientific, Waltham, MA, USA) at 5 m/s for 30 s with 1.4 mm ceramic beads (15-340-153, Fisher Scientific, Waltham, MA, USA). HH37 and older lower jaws were homogenized at 5 m/s for 60 s with 2.8 mm ceramic beads (15-340-154, Fisher Scientific, Waltham, MA, USA). Following purification of total RNA, residual genomic DNA was removed using TURBO DNA-free Kit (AM1907, Invitrogen, Carlsbad, CA, USA). DNased RNA was reverse-transcribed using iSCRIPT (1708841, Bio-Rad). Gene expression was analyzed by quantitative PCR (qPCR) with iQ SYBR Green Supermix (1708882, Bio-Rad, Hercules, CA, USA) and normalized to 18S rRNA following previously published protocols (Dole et al., 2015; Smith et al., 2016). Primers were designed using Geneious Prime (Version 2020.2.4, Auckland, New Zealand) to amplify conserved regions among chick, quail, and duck for *Tgfβr1*, *Ocn*, *Col1a1*, *Mmp2*, and *Mmp13*. Criteria for primer design included limiting primers to 20 bp in length, amplifying regions of ∼150 bp, using an annealing temperature of 60°C, keeping GC content around 50%, minimizing self-complementarity (i.e., primer-dimers), and amplifying regions that span exon-exon junctions. To account for alternative efficiencies of primer binding between species, data were normalized using serial dilutions of pooled cDNA and a standard curve method (Ealba and Schneider, 2013; Dole et al., 2015; Smith et al., 2016). To assay for expression of the eight *Runx2* isoforms, we designed primers to domains containing promoter 1 (P1), promoter 2 (P2), alternatively spliced exon 5 (AS+ or AS-), C-terminus 1 (C1), and C-terminus 2 (C2). Primer sets are listed in **Supplemental Table 1**. Each sample was assayed in technical duplicate.

### Western Blots

Lower jaws were lysed with 1X RIPA lysis buffer (20-188, MilliporeSigma, Burlington, MA, USA) containing Halt protease inhibitors (78430, Thermo Fisher Scientific, Waltham, MA, USA). A BCA assay (23225, Thermo Fisher Scientific, Waltham, MA, USA) was performed to quantify protein using a SpectraMax M5 microplate reader (Molecular Devices, San Jose, CA, USA). 40 µg of protein was electrophoresed on a 10% SDS polyacrylamide gel as previously described (Smith et al., 2016). Proteins were transferred to an Immobilon-P PVDF membrane (IPVH00010, MilliporeSigma, Burlington, MA, USA). Membranes were probed with 1:1000 rabbit anti-chick RUNX2 antibody (ab23981, Abcam, Cambridge, UK), 1:4000 mouse anti-chick β-actin antibody (NB600-501, Novus Biologicals, Littleton, CO, USA), 1:15000 goat anti-rabbit IRDye 800CW (925-32211, LI-COR, Lincoln, NE, USA), and 1:15000 donkey anti-mouse IRDye 680RD antibody (925-68072, LI-COR, Lincoln, NE, USA). Fluorescent signal was detected using the Odyssey Imaging System (LI-COR, Lincoln, NE, USA,). Quantifications of protein bands were performed using Image Studio Lite. Protein levels were normalized to β-actin.

### RNA Isoform Sequencing, Data Alignment, and Normalization

cDNA was synthesized using NEBNext Single Cell/Low Input cDNA Synthesis & Amplification Module (E6421, NEB, USA) following PacBio Iso-Seq template preparation guidelines. cDNA was barcoded using the PCR barcoding expansion 1-96 kit (EXP-PBC096, Oxford Nanopore Technologies, Oxford, UK). cDNA samples were pooled and a sequencing library was prepared using the ligation sequencing kit (Cat # SQK-LSK110, ONT, UK). 50 fmol of the final library was loaded onto each of the two PromethION flowcells (v R9.4.1) and the run was performed for 72 hours. Basecalling and de-multiplexing was performed live on the PromethION compute module with basecaller version ont-guppy-for-minkow 4.0.11.

To determine the number of *Runx2* isoforms, Nanopore sequenced reads were aligned to the runt domain of each respective species using minimap2 (Li, 2018) in Geneious Prime (Prime, 2019). Aligned reads were then aligned to *Runx2* mRNA sequence using Geneious Mapper at the highest sensitivity with the maximum gap size increased to 1000. Aligned reads that did not unambiguously include either P1 or P2 sequence, presence or absence of exon 5, and either C1 or C2 sequence were manually removed. The remaining aligned reads were then counted to determine the number reads for each *Runx2* isoform and the proportion relative to other isoforms. Given the unique challenges of normalizing RNA-seq data among different species (Zhou et al., 2019), we assessed relative expression by normalizing the number of reads for each *Runx2* isoform by total reads per sample.

### Jaw Explants and Cell Culture

For *ex vivo* experiments, HH34 lower jaws were dissected and cultured in 6 well transwell inserts (10769-192, VWR, Radnor, PA, USA) in DMEM complete media. Lower jaws were placed on a 0.45 µm membrane filter (HAWP01300, MilliporeSigma, Burlington, MA, USA) within the transwell inserts. For TGFβ treatments, lower jaws were cultured for 24 hours in complete media, switched to media supplemented with 1% FBS and 1X penicillin-streptomycin, and treated with 25 ng/ml recombinant (r) human TGFβ1 derived from HEK293 cells (100-21, PeproTech, Rocky Hill, NJ, USA) for 24 hours.

For *in vitro* experiments, an embryonic chick fibroblast cell line (DF-1, CRL-12203, ATCC, Manassas, VA, USA) and an embryonic duck fibroblast cell line (CCL-141, ATCC, Manassas, VA, USA) were cultured in complete media, Dulbecco’s Modified Eagle’s Medium (DMEM, 10-013-CV, Corning, Corning, NY, USA) for chick fibroblasts (i.e., DF-1 cells) or MEMα (A10490-01, Thermo Fisher Scientific, Waltham, MA, USA) for duck fibroblasts (i.e., CCL-141 cells) supplemented with 10% Fetal bovine serum (FBS, 97068-085, Lot# 283K18, VWR, Radnor, PA, USA) and 1X penicillin-streptomycin (15140122, Thermo Fisher Scientific, Waltham, MA, USA).

### Generation of Runx2 Isoform Constructs

To generate *Runx2* overexpression constructs, full length cDNA was synthesized using Maxima H-reverse transcriptase (K1651, Thermo Fisher Scientific, Waltham, MA, USA) following the manufacturer’s protocol with 2 μg of total HH37 chick, quail, or duck jaw RNA and 100 pmol of d(T)20 VN primer. The cDNA synthesis reaction was carried out at 50°C for 30 min, 55°C for 10 min, 60°C for 10 min, 65°C for 10 min, and 85°C for 5 min. Full length *Runx2* was amplified by PCR using Q5 Hot Start High-Fidelity DNA Polymerase (M0493L, NEB, Ipswich, MA, USA) and cloned using CloneJET PCR Cloning Kit (K1231, Thermo Fisher Scientific, Waltham, MA, USA). Following confirmation of obtaining full length *Runx2* by Sanger sequencing, *Runx2* was cloned into the pPIDNB inducible-promoter system (Chu et al., 2020), which was digested with AflII (R0520S, NEB, Ipswich, MA, USA) and PstI (R3140S, NEB, Ipswich, MA, USA), using NEBuilder HiFi DNA Assembly Master Mix. pPIDNB contains 1) a piggyBac transposon (Lacoste et al., 2009; Yusa et al., 2011; Yusa, 2015), which allows for stable integration of the construct into a host genome when combined with a pNano-hyPBase plasmid; 2) a constitutively active mNeongreen (GFP) (Shaner et al., 2013), which serves as a reporter for transfection or electroporation efficiency; and 3) a doxycycline (dox)-inducible (Gossen et al., 1995; Loew et al., 2010; Heinz et al., 2011) mScarlet-I (RFP) (Bindels et al., 2017), which serves as a reporter for overexpression of the gene of interest (Chu et al., 2020). All constructs were verified by sequencing and midi-prepped for electroporation using PureLink Fast Low-Endotoxin Midi Kit (A36227, Invitrogen, Carlsbad, CA, USA).

### Electroporation

*In ovo* electroporations were performed as described previously (Chu et al., 2020). Briefly, approximately 0.4 µl of a solution of Fast Green dye plus pEPIC1.1-*Runx2* 1A1 or 2B2 at 3 µg/µl and pNano-hyPBase at 1 µg/µl were mixed, and approximately 0.05 ul was injected with a Pneumatic PicoPump (PV830, World Precision Instruments, Sarasota, FL, USA) into HH8.5 quail and duck anterior neural tubes using thin wall borosilicate glass micropipettes (B100-75-10, O.D. 1.0 mm, I.D. 0.75 mm, Sutter Instrument) pulled on a micropipette puller (P-97 Flaming/Brown, Sutter Instrument, Novato, CA, USA). Platinum electrodes were positioned on each side of the area pellucida, centered at the midbrain-hindbrain boundary and along the neural folds, and three square pulses (1-ms long, 50 volt, with 50 ms spaces), followed by five square pulses (50-ms long, 10 volt, with 50 ms spaces), were administered (CUY21EDITII Next Generation Electroporator, BEX CO, Ltd, Tokyo, Japan) as done previously to allow unilateral entry of the DNA into the presumptive NCM destined for the mandibular arch (Creuzet et al., 2002; Krull, 2004; McLennan and Kulesa, 2007; Hall et al., 2014). The contralateral (un-electroporated) side served as an internal control. Embryos were allowed to develop until HH34 and then were treated *in ovo* with a single dose of 3.75 µg of doxycycline (dox) hyclate (446060250, Acros Organics, Fair Lawn, NJ, USA) in 750 µl of Hanks’ balanced salt solution (14170120, Thermo Fisher Scientific, Waltham, MA, USA) for duck and 0.75 µg of dox in 200 µl of Hanks’ balanced salt solution for quail. Electroporation efficiency and extent of overexpression were evaluated at the time of embryo collection by detecting GFP and RFP on a stereo dissecting microscope under epifluorescent illumination (MZFLIII-TS, Leica Microsystems, Buffalo Grove, IL, USA).

### Luciferase Assay

p6OSE2-Luc was obtained from Dr. Tamara Alliston (UCSF), but was originally generated by Dr. Gerard Karsenty and colleagues (Ducy et al., 1997). p6OSE2-Luc contains 6 tandem copies of the RUNX2 consensus binding motif (i.e., AACCAC) upstream of a luciferase coding sequence in the pGL2 backbone. DF-1 cells were plated at 65,000 cells/cm^2^ in 24-well plates (353047, Corning, Corning, NY, USA) or 250,000 cells/cm^2^ in 6-well plates (354721, Corning, Corning, NY, USA). Cells were transfected in each well using 1.5 μl Lipofectamine 3000 (L3000008, Invitrogen, Carlsbad, CA, USA), 1.5 µl P3000 reagent, 150 ng of β-Gal transfection efficiency control construct, and 400 ng of *Runx2* overexpression construct, and 800 ng of p6OSE2-Luc per well or 750 ng of *Mmp13* promoter-Luc per well. Cells were transfected for 18 hours, then recovered in complete media for 8 hours, and then serum deprived in DMEM without FBS for 18 hours. Cells transfected with pPIDNB-*Runx2* were treated with a final concentration of 100 ng/ml dox in DMEM or MEMα without FBS for 24 hours. Cells were lysed in 1X lysis buffer (E1531, Promega, Madison, WI, USA) and analyzed for luciferase activity using beetle luciferin (E1602, Promega, Madison, WI, USA) and Coenzyme A (J13787MF, ThermoFisher Scientific, Waltham, MA, USA) normalized to β-galactosidase activity using Galacto-Star β-Galactosidase Reporter (T1012, Invitrogen, Carlsbad, CA, USA) as described (Chen et al., 2012b). Luminescence was measured using a Spectramax M5 luminometer. At least two preparations of each DNA construct were tested for over-expression experiments.

### Mmp13 Promoter Sequencing and Generation of Mmp13 Promoter Constructs

Previously, we identified RUNX2 binding sites in the *Mmp13* promoter (Smith et al., 2020) using the JASPAR 2020 database, which contains transcription factor-binding profiles stored as position frequency matrices (Fornes et al., 2020). Briefly, to map transcription factor-binding sites onto the *Mmp13* promoter sequences of chick, quail, and duck we used TFBSTools (Tan and Lenhard, 2016), which is an R package (Team, 2013). For the proximal region of the *Mmp13* promoter (*i.e.*, −184 bp for chick and quail, and −181 bp for duck) all vertebrate transcription factors were included in the analysis. Position frequency matrices were converted to position weighted matrices by setting pseudocounts to 0.8 (Nishida et al., 2009), and background frequencies of nucleotides to 0.25. The minimum threshold score was set to 95% for the −184/181 bp promoter region and 90% for the −2 kb promoter region.

In a previous study we generated luciferase constructs containing regions of the *Mmp13* promoter (Smith et al., 2020). Briefly, *Mmp13* promoter sequences for chick, quail, and duck were amplified by PCR using Q5 Hot Start High-Fidelity DNA Polymerase (M0493L, NEB, Ipswich, MA, USA). pGL3 was digested with HindIII-HF (R3104S, NEB, Ipswich, MA, USA) and XhoI (R0146S, NEB, Ipswich, MA, USA). The amplified *Mmp13* promoter sequences and digested pGL3 were purified using GeneJET PCR Purification Kit and cloned using NEBuilder HiFi DNA Assembly Master Mix (E2621L, NEB, Ipswich, MA, USA). Mutations in *Mmp13* promoter SNPs were generated through site-specific mutagenesis PCR with the mutations in the primers (Ho et al., 1989). All constructs were verified by sequencing and midi-prepped for transfection using PureLink Fast Low-Endotoxin Midi Kit (A36227, Invitrogen, Carlsbad, CA, USA).

### Statistics and Image Processing

Data are represented as a mean ± standard error of the mean (SEM). Statistical significance was determined through two-tailed ANOVA adjusted for multiple comparisons using the Bonferoni (Prism, Version 9.1.0, GraphPad, San Diego, CA). A Student’s t-test was used for comparisons between *Runx2* isoform overexpression and contralateral controls. A Benjamini–Hochberg procedure was used for luciferase experiments comparing *Runx2* isoform overexpression to empty vector controls. For *in ovo* data, “n” refers to the total number of embryos analyzed per group. For *in vitro* data, “n” refers to the total number of individual wells analyzed per group, with each experiment replicated at least three times. In all figures, p ≤ 0.05 is considered statistically significant, although some statistical comparisons reached significance below p ≤ 0.01, p ≤ 0.001, or p ≤ 0.0001 where noted. Formal power analyses were not conducted, and group size “n” is denoted in the figure legends. Images of tissue sections and embryos were adjusted in Adobe Photoshop 2020 (version 21.2.2) to normalize for exposure, brightness, contrast, saturation, and color balance across samples. Figures were assembled in Adobe Illustrator 2020 (Version 24.2.3).

## RESULTS AND DISCUSSION

### The Overall Structure of Runx2 is Highly Conserved Across Vertebrates

We compiled a list of annotated *Runx2* sequences on NCBI and find 226 species. We then compared these with NCBI sequences for human, mouse, chick, and zebrafish, as well as sequences for domains of *Runx2* to determine if the NCBI sequences included the complete *Runx2* gene. Around 55% (123 out of 226) of the species examined with an annotated *Runx2* gene on NCBI appear to be incomplete. Nearly all of the incomplete sequences are missing N-terminal exons and some sequences only contain a single C-terminal exon. We expect that most if not all of the incomplete sequences in NCBI are related to sequencing, assembly, and bioinformatics issues rather than representing actual *Runx2* gene evolution. Most *Runx2* truncations appear to occur near the QA repeat, which may reflect the difficulty of sequencing and generating contigs of repetitive sequences. We also find that most of the incomplete sequences contain gaps or highly repetitive sequences directly upstream of the truncated *Runx2* sequences. Additionally, every sequence analyzed that does not contain either P1 or P2 promoter sequences also lacks a QA repeat. Moreover, the duck and quail *Runx2* sequences on NCBI appear to be incomplete and inaccurate. The duck *Runx2* sequence on NCBI lacks both P1 and P2 promoters and the QA repeat. By running 5’ RACE reactions on duck mRNA, we confirmed the presence of both P1 and P2 promoters as well as the QA repeat in the duck *Runx2* gene. The quail *Runx2* sequence on NCBI lacks the P1 promoter and also contains an alternative noncanonical promoter. By running 5’ RACE reactions on quail cDNA we confirmed the presence of the P1 and P2 promoters but did not detect the presence of an alternative noncanonical promoter.

We compared full length RUNX2 for chondrichthyian (n = 1), actinopterygian (n = 25), sarcopterygian (n = 1), amphibian (n = 5), reptilian (n = 11), avian (n = 13), and mammalian (n = 46) taxa as listed in **Supplemental Table 2**. We find that most of the RUNX2 domains are conserved as shown in a similarity matrix of AA sequences (**Figure 1B**). However, the QA domain encoded by exon 2 and C-terminus 2 (C2) encoded by exon 9 are so variable that we did not include them in the similarity matrix. We find that the structure and splice points for all exons of *Runx2* are highly conserved among the taxa analyzed. Exon 1 encodes for 19 AA, exon 3 encodes for 52 AA, exon 4 encodes for 35 AA, exon 5 encodes for 58 AA (except for actinopterygians, which have 59 AA), exon 6 encodes for 54 AA (except for chondrichtyians, which have 55 AA), and exon 8 encodes for 159 to 169 AA. Exon 7, which is specific to mammals, encodes for 22 AA. In terms of pairwise identity of AA for individual domains across vertebrates, the most conserved is the RHD, which is encoded by part of exon 2 (with 98.3% identity), all of exon 3 (with 97.8% identity), and all of exon 4 (with 98.9% identity). Conservation of the RHD is even higher within amniotes and within each major clade. This is not surprising given that the RHD is an ancient 128 AA DNA binding domain that arose early in metazoan evolution and defines the RUNX protein family (Kagoshima et al., 1993; Rennert et al., 2003; Levanon and Groner, 2004; Coffman, 2009; Robertson et al., 2009; Chuang et al., 2013; Ito et al., 2015; Mevel et al., 2019). Interestingly in reptiles (including birds), we find that the RHD is modified by one AA, which goes from phenylalanine (F) to leucine (L). Overall, amniotes have higher conservation than vertebrates as a whole, and birds have the highest levels of conservation with seven domains in the similarity matrix showing 100% pairwise identity of AA, and two domains (alternatively spliced exon 5 and exon 8, which encodes C1) showing more than 99% identity. Consensus sequences for the major vertebrate lineages are shown in **Supplemental Table 3**.

### RUNX2 Has Undergone Key Evolutionary Changes in Four Domains

Using parsimony, we mapped the shared derived changes in RUNX2 structure on the major vertebrate lineages. We find that the actinopterygian, amphibian, reptilian, and mammalian taxa each cluster into distinct monophyletic groups (**Figure 2A**). RUNX2 shows the most divergence in the actinopterygian lineage, which can be split into two branches (not shown) with one cluster comprised entirely of teleosts and the other containing four species that do not form a natural group (i.e., spotted gar, red-bellied piranha, channel catfish, and one of two *Runx2* copies found in zebrafish). Perhaps the whole genome duplication that occurred in the teleost lineage (Glasauer and Neuhauss, 2014) has led to greater evolutionary divergence in *Runx2* sequence. Only a single copy of *Runx2* was identified for the teleosts we examined except for zebrafish, which is known to retain both copies (Flores et al., 2004; Eames et al., 2012). Therefore, one copy of *Runx2* was either lost among the majority of teleosts, which has been the common fate for many genes following the teleost-specific genome duplication (Pasquier et al., 2017), or simply cannot be identified at present using the available bioinformatic data. Overall, our comparative analysis of RUNX2 reveals key evolutionary events that occurred primarily in four domains: a domain containing QA repeats encoded by exon 2, a domain containing a pentapeptide at the start of the canonical C-terminus encoded by exon 8, a domain consisting of a novel C-terminus encoded by exon 9 in amniotes, and a domain encoded by an alternatively spliced exon 7 in mammals (**Figure 2A**).

**Figure 2.**
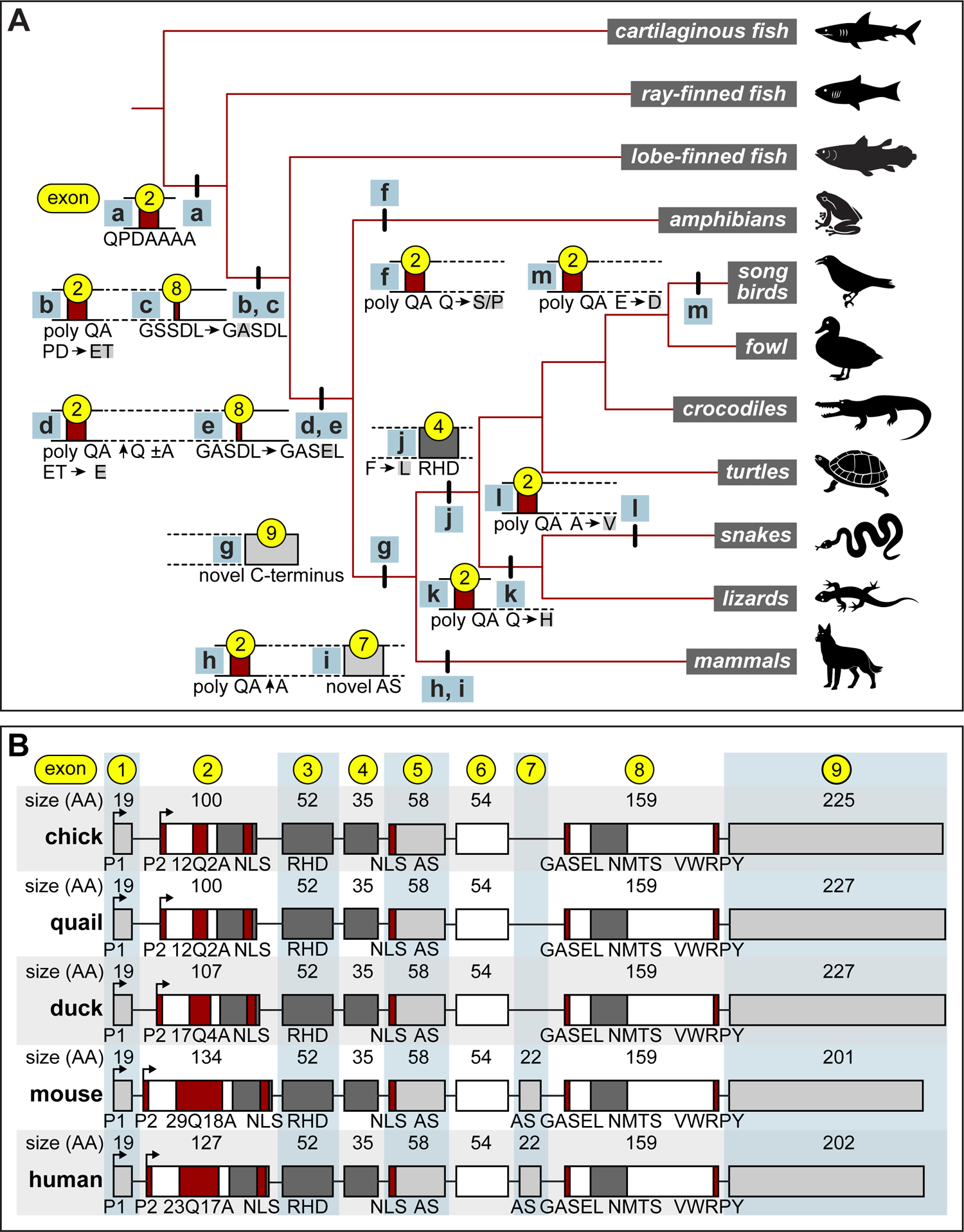
Evolution of RUNX2 domains across vertebrates. **A)** Shared derived changes in RUNX2 structure can be mapped on the major vertebrate lineages. Based on parsimony, the common ancestor of actinopterygian (ray-finned fish) and sarcopterygians (lobe-finned fish) likely had a QA domain with some poly A repeats encoded by exon 2 as indicated by the “a” hash mark at the base of the ingroup from chondrichthyans (“cartilaginous fish”). The common ancestor of sarcopterygians and tetrapods (amphibians, reptiles (including birds), and mammals) likely had a QA domain with an additional Q near the poly A repeats encoded by exon 2 and a change in the GSSDL pentapeptide to GASDL encoded by exon 8 as indicated by the “b,c” hash mark. The common ancestor of amphibians and amniotes (reptiles and mammals) likely had an expansion in the length of the poly Q repeat and the pentapeptide GASDL changed to GASEL as indicated by the “d,e” hash mark. In amphibia the poly Q repeat is interspersed with serine, proline, and/or histidine (as indicated by the “f” hash mark) while amniotes primarily contain a homomeric poly Q repeat. The common ancestor of amniotes likely had a novel C-terminus as indicated by the “g” hash mark. The common ancestor of mammals likely had an expansion in the length of the poly A repeat and a novel alternatively spliced exon 7 as indicated by the “h,i” hash mark. In reptiles (including birds), the RHD is modified by one AA, which goes from phenylalanine to leucine (as indicated by the “j” hash mark). In the squamate reptiles (snakes and lizards), the poly Q repeat is interspersed with histidine (as indicated by the “k” hash mark). In snakes, the poly A repeat is interspersed with valines (as indicated by the “l” hash mark). In the passeriform birds (songbirds), a glutamic acid spacer between the poly Q and poly A repeats is changed to aspartic acid as indicated by the “m” hash mark. **B)** Schematic showing the number of amino acids (AA) encoded by each exon of *Runx2* for chick, quail, duck, mouse, and human. *Runx2* contains two alternative promoters (P1 and P2), repeats of glutamine (Q) and alanine (A), which vary in length depending on the taxon; nuclear localization signals (NLS); a runt homology domain (RHD); an alternatively spliced (AS) exon 5 in all taxa as well exon 7, which is specific to mammals; and two alternative C-termini (C1 and C2). C1 (exon 8) encodes pentapeptide domains (GASEL and VWRPY) and a nuclear matrix targeting signal (NMTS). C2 (exon 9) is highly variable.

For the first domain, we hypothesize that the common ancestor of actinopterygians (ray-finned fish) and sarcopterygians (lobe-finned fish) likely evolved a few poly A repeats encoded by exon 2 and this distinguishes them from chondrichthyans (cartilaginous fish). The common ancestor of sarcopterygians and tetrapods (amphibians, reptiles (including birds), and mammals) likely had a QA domain with an additional Q near the poly A repeats. The common ancestor of amphibians and amniotes (reptiles, birds, and mammals) likely had an expansion in the length of the poly Q repeat. We find that in amphibia the poly Q repeat is interspersed with serine, proline, and/or histidine while amniotes primarily contain a homomeric poly Q repeat. In the squamate reptiles (snakes and lizards), the poly Q repeat is interspersed with histidine, and further in snakes, the poly A repeat is interspersed with valines. In the passerine birds (songbirds), a glutamic acid spacer between the poly Q and poly A repeats is changed to aspartic acid. The common ancestor of mammals likely had an expansion in the length of the poly A repeat. While the functional significance of these evolutionary changes in AA composition is not well understood, the expansion of the QA domain corresponds with the emergence of novel traits in vertebrates (Newton and Pask, 2020) and further, the ratios of Q to A appear to affect RUNX2 transcriptional activity and correlate with facial length in some mammals (Fondon and Garner, 2004; Sears et al., 2007; Morrison et al., 2013; Mastushita et al., 2015; Ritzman et al., 2017; Ferraz et al., 2018) but not others (Pointer et al., 2012; Newton et al., 2017).

With regard to the second domain, which contains a pentapeptide at the start of the canonical C-terminus encoded by exon 8, the common ancestor of sarcopterygians and tetrapods likely underwent a change from GSSDL to GASDL. Then GASDL changed to GASEL in the common ancestor of amphibians and amniotes. In previous studies, the GASEL pentapeptide motif has been shown to be an activating domain of RUNX2 and is sufficient to drive increased transcription (Thirunavukkarasu et al., 1998). None of the fish we examined contain the exact GASEL sequence; however, there is a similar and predominant actinopterygian motif of GSSDL, which is found in 18 of 25 taxa analyzed. Sequences from two medaka and one copy of the zebrafish RUNX2 contain a GSTDL motif while the other copy of the zebrafish RUNX2 contains GTSEL. The channel catfish and red-bellied piranha have a GTSDL motif. Surprisingly, the shark, spotted gar, and coelacanth all contain the same pentapeptide sequence of GASDL, which is only a single AA different than GASEL. With the exception of one copy of the zebrafish *Runx2* and the GASDL, the other actinopterygian sequences are two AA different than GASEL. What remains unknown is whether GASDL is the ancestral motif or if this arose independently several times. One amphioxus (Stricker et al., 2003), one hagfish (Hecht et al., 2008), and three Japanese lamprey *Runx* genes have been identified; however, which lamprey *Runx* genes match with the gnathostome *Runx* genes or if they even match on a one-to-one basis is unclear (Nah et al., 2014). The lamprey *Runx* genes contain ASSEL, GGSEL, and DLQGV at the motif site, all of which are at least two AA different than GASDL indicating that the GASDL sequence may not be ancestral. All of the amphibian, reptilian, avian, and mammalian sequences contain the exact same GASEL pentapeptide motif.

In relation to the third domain, the common ancestor of amniotes likely evolved a novel C-terminus. This additional alternatively spliced C-terminal domain was originally discovered in mouse, humans, and rat, and designated as exon 6.1 (Terry et al., 2004). We label this domain as exon 9 and find homologous sequence only among amniotes (**Figure 2B**). Nucleotide and protein BLAST queries using chicken and mouse exon 9 sequences against the NCBI database of actinopterygian and amphibian sequences did not result in any hits for *Runx2*. Compared to the rest of the RUNX2 sequence (with the possible exception of the QA repeat), C2, which is encoded by exon 9, is the least conserved (**Supplemental Table 4**). For example, chicken and mouse RUNX2 AA sequences are 96.4% identical when excluding the region of exon 2 that encodes the QA repeat, exon 7, and exon 9, but are only 36.3% identical when comparing the AA sequences encoded by exon 9. Moreover, chicken and duck are 99.6% identical when excluding the region of exon 2 that encodes the QA repeat and exon 9 but are only 79.7% identical when comparing the AA sequences encoded by exon 9.

For the fourth domain, the common ancestor of mammals likely evolved a novel alternatively spliced exon 7. All of the mammal *Runx2* sequences analyzed (n = 46) appear to contain exon 7 but we were unable to find similar sequences after running nucleotide and protein BLAST queries using human or mouse exon 7 sequences against the NCBI database of actinopterygian, amphibian, reptilian, or avian sequences. The alternative splicing of exon 7 in mammals (along with exon 5) is known to cause changes in nuclear localization of RUNX2 protein and has different functional effects on target gene expression (Makita et al., 2008). We further confirmed that exons 5 and 7 are alternatively spliced in the developing jaw primordia of mammals by examining *Runx2* transcripts in a publicly available RNA-seq dataset (Len Pennacchio, 2017) from mouse at embryonic day 13.5 (data not shown). The splicing out of exon 5 in mammals appears to be coincident with the splicing out of exon 7, and the absence of exon 7 has been shown to reduce the activating potential of RUNX2 (Terry et al., 2004; Makita et al., 2008). We have found that exon 5 is also alternatively spliced in birds but whether this is the same for other non-amniote taxa remains unclear. Despite these evolutionary changes in RUNX2 structure, the extent of RUNX2 conservation among amniotes is striking and clearly evidenced by the fact that each exon encodes the exact same number of AA in chick, quail, duck, mouse, and human (**Figure 2B**). We only find major differences in exon 2, which encodes for the QA repeats that vary in length depending on the taxon, and exon 9, which encodes for the alternatively spliced C-terminus.

### Runx2 Expression is Coincident with Bone Deposition and is Species-Specific

Given, the high degree of conservation in RUNX2 structure across vertebrates, we investigated the extent to which developmental changes in spatial and temporal patterns of expression, and/or the deployment of isoforms may contribute to the evolution of the jaw skeleton. We focused on birds that show species-specific differences in jaw size and shape. Our prior work involving transplants between quail and duck embryos has demonstrated that the timing of induction and levels of *Runx2* expression during osteogenesis in the jaw are mediated by NCM (Merrill et al., 2008; Hall et al., 2014).

To confirm that areas of bone deposition co-localize with RUNX2 expression in the lower jaw, we stained the osteoid matrix and assayed for RUNX2 protein using IHC on adjacent sections of HH37 quail (**Figure 3A and 3B**) and duck (**Figure 3C and 3D**). We find that osteoid coincides with RUNX2 protein distribution in proximal regions of the quail (**Figure 3E and 3F**) and duck (**Figure 3G and 3H**) lower jaws such as in the angular bone; and in the more distal dentary bone of the quail (**Figure 3I and 3J**) and duck (**Figure 3K and 3L**). We did not observe any remarkable spatial or specific-specific differences in RUNX2 protein localization between quail and duck.

**Figure 3.**
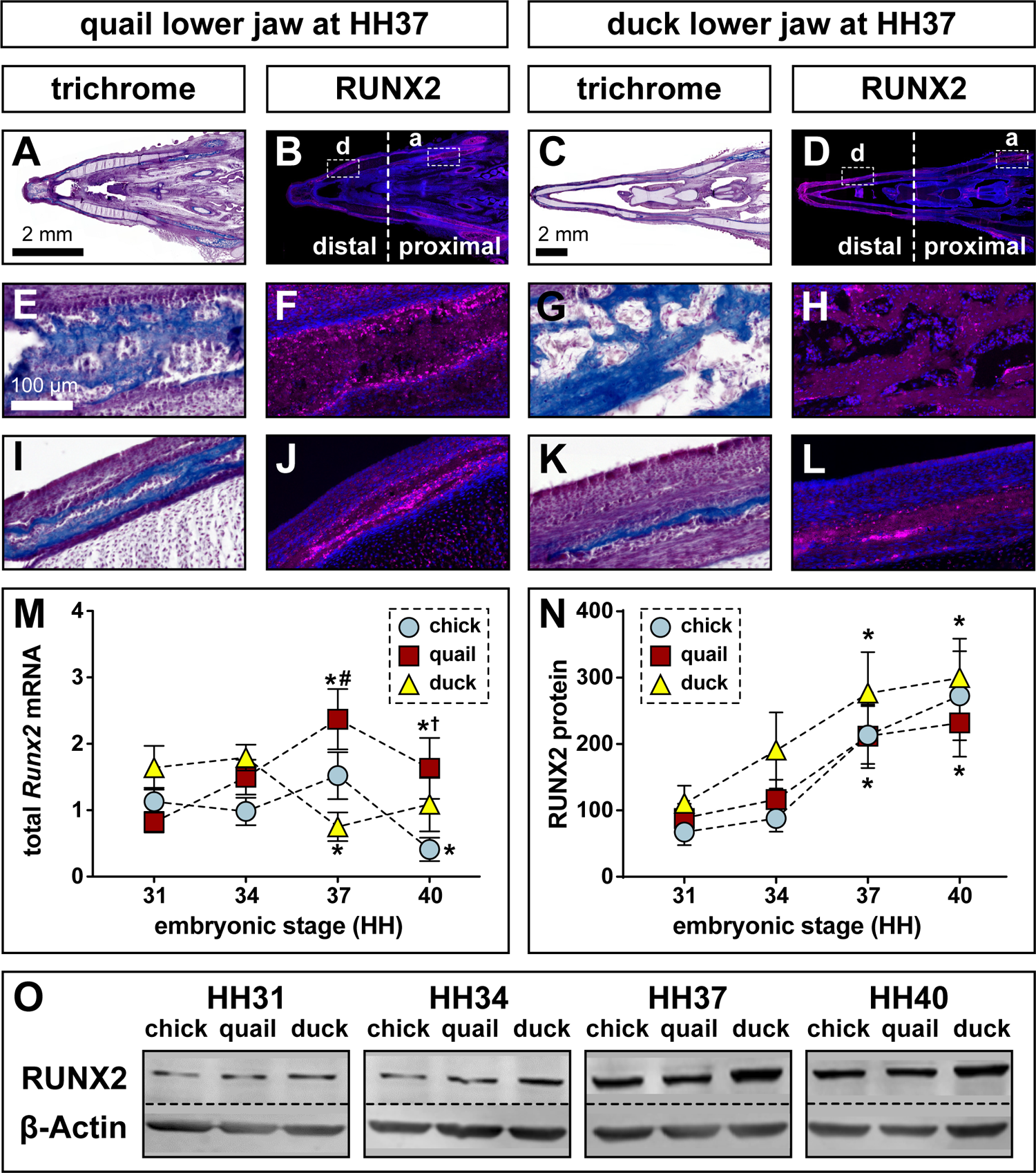
Species-specific expression of RUNX2. **A)** Sections of quail lower jaw stained with Trichrome (osteoid matrix of bone is blue) or **B)** RUNX2 antibody (pink) and Hoechst dye (blue) that stains nuclei at HH37 (n = 4). RUNX2 protein can be detected in the dentary (d) bone in the distal portion of the lower jaw and in the angular (a) bone in the proximal portion. **C)** Sections of duck lower jaw stained with Trichrome or **D)** RUNX2 antibody at HH37 (n = 4). **E)** Higher magnification images of the angular bone stained with Trichrome and **F)** RUNX2 antibody in quail; and **G)** Trichrome and **H)** RUNX2 antibody in duck. I) Higher magnification images of the dentary bone stained with Trichrome and **J)** RUNX2 antibody in quail; and **K)** Trichrome and **L)** RUNX2 antibody in duck. **M)** qPCR analysis of *Runx2* mRNA expression shows a 2.6-fold increase in quail (red square) and a 2.4-fold decrease in duck (yellow triangle) from HH31 to HH37, and quail *Runx2* expression is significantly higher than duck at HH37 (n = 8). There are no significant changes in chick (blue circle) until HH40 when *Runx2* decreases by 2-fold (n = 8). **N)** RUNX2 protein quantification and **O)** Western blots show a 4-fold increase in chick (n = 9) and a 3-fold increase in quail and duck (n = 12) from HH31 to HH37. RUNX2 protein continues to increase at HH40. No significant differences in RUNX2 protein levels are observed among the species at any stage.

However, when we examine *Runx2* mRNA levels among chick, quail, and duck from HH31 to HH40, we do observe species-specific differences in expression when using qPCR (**Figure 3M**). We quantified expression in chick, quail, and duck jaws across several stages of development, and compared expression relative to HH31 for each species. In chick, we observe no significant change in total *Runx2* levels until HH40, where expression decreases by 2.8-fold (p ≤ 0.05). In quail, total *Runx2* expression increases 2.9-fold at HH37 (p ≤ 0.003) and 2-fold HH40 (p ≤ 0.05). At HH37, quail has 3.2-fold higher *Runx2* expression compared to duck (p ≤ 0.0009). In duck, total *Runx2* decreases by 2.2-fold at HH37 (p ≤ 0.05) but increases at HH40 to levels like that observed at HH31. These data are consistent with our prior observations that quail and duck express equivalent levels of *Runx2* at early stages (i.e., HH24 to HH32) but that quail expresses more than twice the amount of *Runx2* at later stages (i.e., HH36 to HH38) of jaw development (Hall et al., 2014).

We quantified RUNX2 protein by western blot (WB) and do not detect any significant species-specific differences but do observe a steady increase in RUNX2 levels from HH31 to HH40 for chick, quail, and duck (**Figure 3N**). WB reveal that RUNX2 protein increases 3.2-fold in chick (p ≤ 0.04), 2.4-fold in quail (p ≤ 0.03), and 1.3-fold in duck (p ≤ 0.05) at HH37; and 4-fold in chick (p ≤ 0.04), 2.6-fold in quail (p ≤ 0.03), and 2.7-fold in duck (p ≤ 0.05) at HH40 with no significant differences observed among the species (**Figure 3O**). Thus, although quail has increased *Runx2* mRNA expression at HH37, we do not observe significant differences at the protein level either by IHC or WB. The differences between what we observe on the mRNA and protein levels suggest there may be post-transcriptional regulation of *Runx2* and/or post-translational regulation of RUNX2 occurring within each species. Such regulation on the gene and protein levels has been well documented for *Runx2* in a range of other contexts (Kim et al., 2003; Shen et al., 2006; Rajgopal et al., 2007; Hassan et al., 2010; Ito et al., 2015; Vimalraj et al., 2015; Narayanan et al., 2019; Kim et al., 2020). Alternatively, the discrepancies among the IHC, WB, and qPCR data may be due to limitations in sensitivity of IHC as non-quantitative and WB as semi-quantitative techniques for measuring expression.

### Runx2 Isoforms are Differentially Expressed in the Developing Jaw

By sequencing *Runx2* mRNA transcripts present in the developing jaw primordia of chick, quail, and duck, we find that alternative splicing produces eight potential avian isoforms (**Figure 4A**). We annotated each isoform to indicate which domains are present such that promoter 1 (P1) = 1, promoter 2 (P2) = 2, alternatively spliced exon 5 included (AS+) = A, alternatively spliced exon 5 excluded (AS-) = B, C-terminus 1 (C1) = 1, or C-terminus 2 (C2) = 2. Thus, the eight avian *Runx2* isoforms are 1A1, 1B1, 1A2, 1B2, 2A1, 2B1, 2A2, 2B2. For example, *Runx2* isoform 1A1 consists of P1, AS+, and C1; and is most distinct from 2B2, which consists of P2, AS-, and C2.

**Figure 4.**
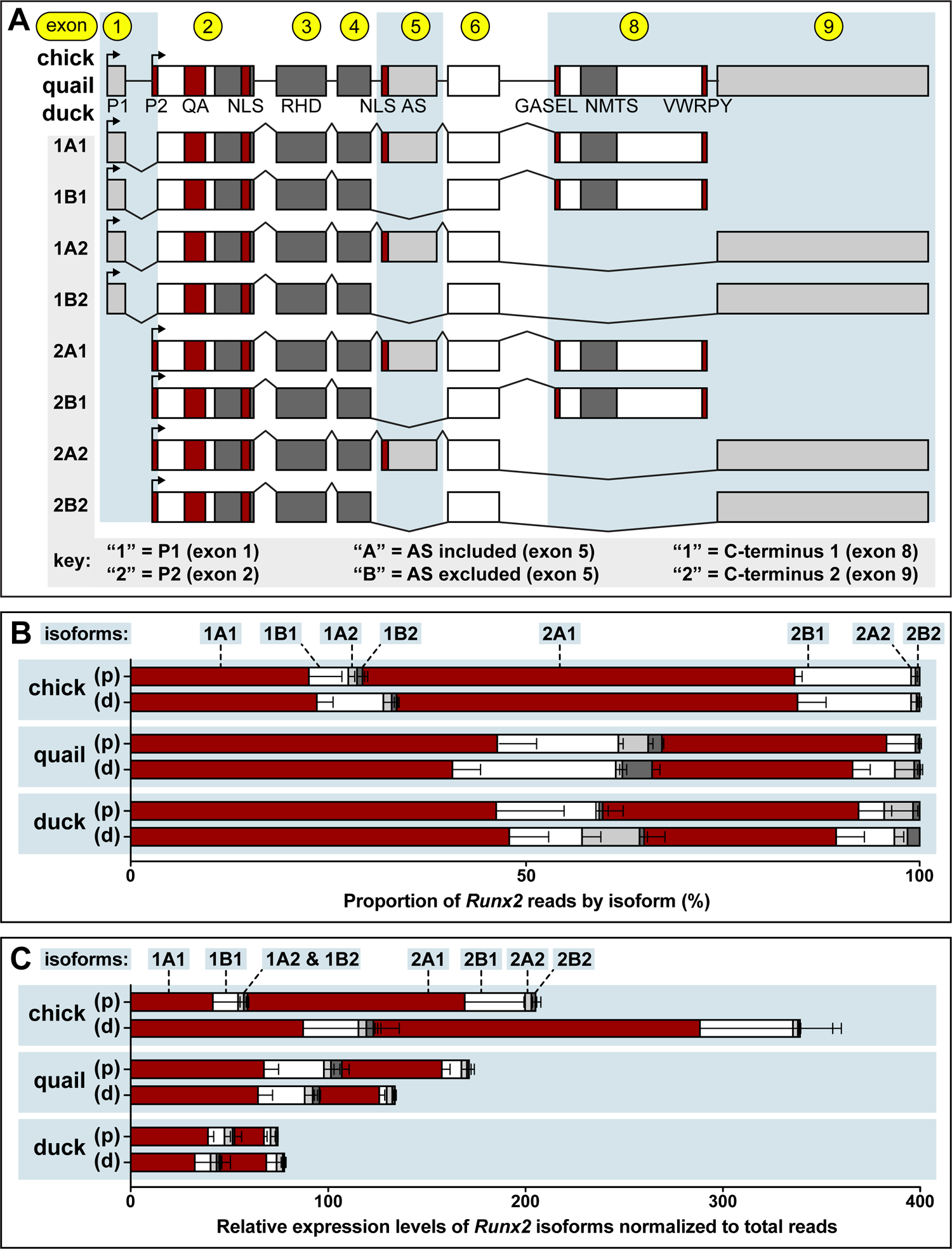
*Runx2* isoforms and their expression in the avian jaw. **A)** Schematic illustrating the eight isoforms of *Runx2* in chick, quail, and duck. Isoforms are annotated based on splicing, indicating presence of promoter 1 (P1) or promoter 2 (P2), the presence (“A)” or absence (“B”) of alternatively spliced (AS) exon 5, and the presence of C-terminus 1 (C1) or C-terminus 2 (C2). Functional domains include the glutamine alanine (QA) repeats, the nuclear localization sequence (NLS), the Runt Homology Domain (RHD), GASEL, NMTS, and VWRPY. **B)** RNA isoform sequencing (Iso-Seq) of chick (n = 2), quail (n = 2), and duck (n = 2) proximal (p) and distal (d) portions of the lower jaws reveals spatial and species-specific variation in the proportions of different isoforms of *Runx2*. Within chick, the *Runx2* 2A1 isoform is most prominently expressed, followed by 1A1, 2B1, and 1B1. 1A2, 2A2, and 2B2 have low or no expression in chick. In quail and duck, the *Runx2* 1A1 isoform is most prominently expressed, followed by 2A1, 1B1, and 2B1. 1A2, 2A2, and 2B2 have low or no expression in quail and duck. **C)** The relative expression of each *Runx2* isoform normalized to total reads shows that overall *Runx2* expression highest in the chick distal portion, followed by chick proximal, quail proximal, quail distal, and duck proximal and distal. 2A1 is most prominently expressed in the chick distal region, followed by 1A1, 2B1, and 1B1. In quail and duck, 1A1 is most prominently expressed, followed by 2A1, 1B1, and 2B1. 1A2, 2A2, and 2B2 have low or no expression in quail and duck. The 1A1, 1B1, and 2A1 are more expressed in quail than duck. Other isoforms are similar among the species. Chick expresses more isoforms utilizing P2, while quail and chick express more isoforms with P1.

To determine the extent to which these eight *Runx2* isoforms are differentially expressed in chick, quail, and duck, we performed an RNA Iso-Seq experiment using HH37 lower jaws, which again is the stage when we observe the greatest amount of species-specific divergence in the levels of *Runx2* mRNA (**Figure 3M**). Also, to assess if these isoforms are spatially regulated in the developing jaw primordia, we divided the jaw samples into distal (containing the dentary bone) and proximal (containing the angular and other bones) portions (**Figure 3B and 3D**). When examining the absolute proportion of *Runx2* isoforms present (**Figure 4B**), we find that for the proximal region of chick lower jaws, 2A1 is the highest at 54%, followed by 1A1 at 22%, 2B1 at 15%, 1B1 at 5%, and the remaining isoforms totaling approximately 4%. Similar percentages are found in distal chick lower jaws, with 2A1 being the highest at 51%, followed by 1A1 at 24%, 2B1 at 14%, 1B1 at 8%, and the remaining isoforms totaling approximately 3%. In contrast, for proximal quail lower jaws, 1A1 is the highest at 46%, followed by 2A1 at 28%, 1B1 at 15%, 1A2 at 3.9%, and the remaining isoforms totaling approximately 7%. Similar percentages are found in distal quail lower jaws, with 1A1 being the highest at 41%, followed by 2A1 at 25%, 1B1 at 20%, 1B2 at 3.8%, and the remaining isoforms totaling approximately 10%. We observe similar results for proximal duck lower jaws, with 1A1 being the highest at 46%, followed by 2A1 at 32%, and 1B1 at 12%, and the remaining isoforms totaling approximately 10%. Similar percentages are found in the distal duck lower jaws for 1A1 at 49% and 2A1 at 25%. However, duck 1B1 is at 9% compared to quail 1B1 at 20%, and duck 1B2 is at 7% compared to quail 1B2 at 2.8%. The remaining isoforms in duck total approximately 10%. Thus, in terms of absolute proportions, more than half of the *Runx2* expressed in the chick lower jaw (both proximal and distal regions) is the 2A1 isoform, whereas the predominant isoform in quail and duck lower jaws (both proximal and distal regions) is the 1A1 isoform. Also, we find that in the distal lower jaw, 1B1 and 1B2 isoforms in quail are inversely proportional to 1B1 and 1B2 isoforms in duck.

To determine if the relative expression of *Runx2* isoforms varies by spatial distribution, and/or by species we normalized the number of reads for each *Runx2* isoform by total reads per sample. We find that in terms of overall relative expression, chick distal lower jaws show the highest levels of *Runx2*, followed by chick proximal, quail proximal, quail distal, duck distal, and duck proximal (**Figure 4C**). We do observe some differences in spatial expression of *Runx2* isoforms between proximal and distal lower jaws. For example, chick has 2.1 times more 1A1 and 1B1; and around 1.7 times more 2A1 and 2B1 in the distal lower jaw than in the proximal lower jaw. When comparing expression among species, we observe that chick 2A1 and 1A1 show the highest levels of expression. We observe higher expression of 2B1 in chick than quail but similar expression for the other isoforms. We observe higher expression of 1A1, 1B1, and 2A1 in quail than duck but similar expression for the other isoforms. Overall, we observe that at HH37, chick lower jaws have the highest *Runx2* isoform expression with 2A1, 1A1, and 2B1 being the most prevalent, followed by quail where 1A1, 2A1, and 1B1 are most prevalent, and with duck having the lowest expression with 1A1, 2A1, and 1B1 being the most prevalent isoforms. Chick deploy more *Runx2* isoforms utilizing P2 while quail and duck deploy isoforms utilizing P1. This species-specific and spatial distribution of *Runx2* isoforms that we observe could potentially provide local developmental control over the regulation of target genes involved in the deposition and resorption of bone and lead to differential growth of the jaw skeleton. Similarly, other studies have proposed that distinct zones of bone remodeling can direct the size and shape of the developing jaw (Enlow et al., 1975; Moore, 1981; Radlanski and Klarkowski, 2001; Radlanski et al., 2004; Ealba et al., 2015).

To assess the extent to which *Runx2* isoforms change over developmental time and to determine if there are species-specific trends in the deployment of certain functional domains, we designed qPCR primers to amplify all isoforms utilizing the P1 domain (i.e., 1A1, 1A2, 1B1, 1B2), P2 domain (i.e., 2A1, 2A2, 2B1, 2B2), AS+ domain (i.e., 1A1, 1A2, 2A1, 2A2), AS-domain (i.e., 1B1, 1B2, 2B1, 2B2), C1 domain (i.e., 1A1, 1B1, 2A1, 2B1), and C2 domain (i.e., 1A2, 1B2, 2A2, 2B2) in the jaw primordia of chick, quail, and duck, at HH31, HH34, HH37, and HH40. Our qPCR analyses reveal that expression of isoforms utilizing the P1 domain increases 3.3-fold in chick (p ≤ 0.04) and 3.5-fold in quail (p ≤ 0.05) by HH37 but decreases in both species by HH40 (**Figure 5A**). P1 expression does not change over time in duck but is significantly higher compared to chick (p ≤ 0.02) and quail (p ≤ 0.008) at HH34. Expression of isoforms utilizing the P2 domain significantly decreases 5.5-fold in chick by HH40 (p ≤ 0.003). P2 expression increases 2-fold in quail by HH37 (p ≤ 0.04), while P2 expression does not change in duck (**Figure 5B**). No significant differences in P2 expression are observed among species. Expression of isoforms containing the AS+ domain is constant in chick until HH40 (**Figure 5C**) when there is a 5-fold decrease (p ≤ 0.03). AS+ increases 3-fold in quail by HH37 (p ≤ 0.002) and is significantly higher than that observed in chick (p ≤ 0.0003) and duck (p ≤ 0.0001). Expression of isoforms defined by the AS-domain increases 2-fold in chick by HH37 (p ≤ 0.05) (**Figure 5D**). AS-expression increases 2.5-fold in quail by HH37 (p ≤ 0.03) and is significantly higher than that observed in duck (p ≤ 0.04). AS-expression is significantly higher in duck, compared to chick (p ≤ 0.03) and quail (p ≤ 0.01) at HH31, but decreases 2.3-fold by HH37 (p ≤ 0.05) and 2.8-fold by HH40 (p ≤ 0.03). Expression of isoforms containing the C1 domain remains constant among all three species, although expression is significantly higher in duck than quail at HH31 (p ≤ 0.002) (**Figure 5E**). Expression of isoforms containing the C2 domain remains constant among all three species, although expression is significantly higher in duck than chick (p ≤ 0.02) and quail (p ≤ 0.006) at HH31 (**Figure 5F**). Overall, we observe changes in *Runx2* domain expression during embryonic development of chick, quail, and duck. Specifically in quail at HH37, we observe an increase in P1, P2, AS+, and AS-expression, and observe a higher expression of AS+ and AS-in quail compared to duck at HH37. The change in *Runx2* isoform expression during development could result in differential transcriptional activity and regulation of target genes, which could lead to species-specific regulation of phenotypic outcomes. To investigate mechanisms through which the differential expression of *Runx2* isoforms could be regulated in the context of avian jaw development, we interrogated the potential role of TGFβ signaling.

**Figure 5.**
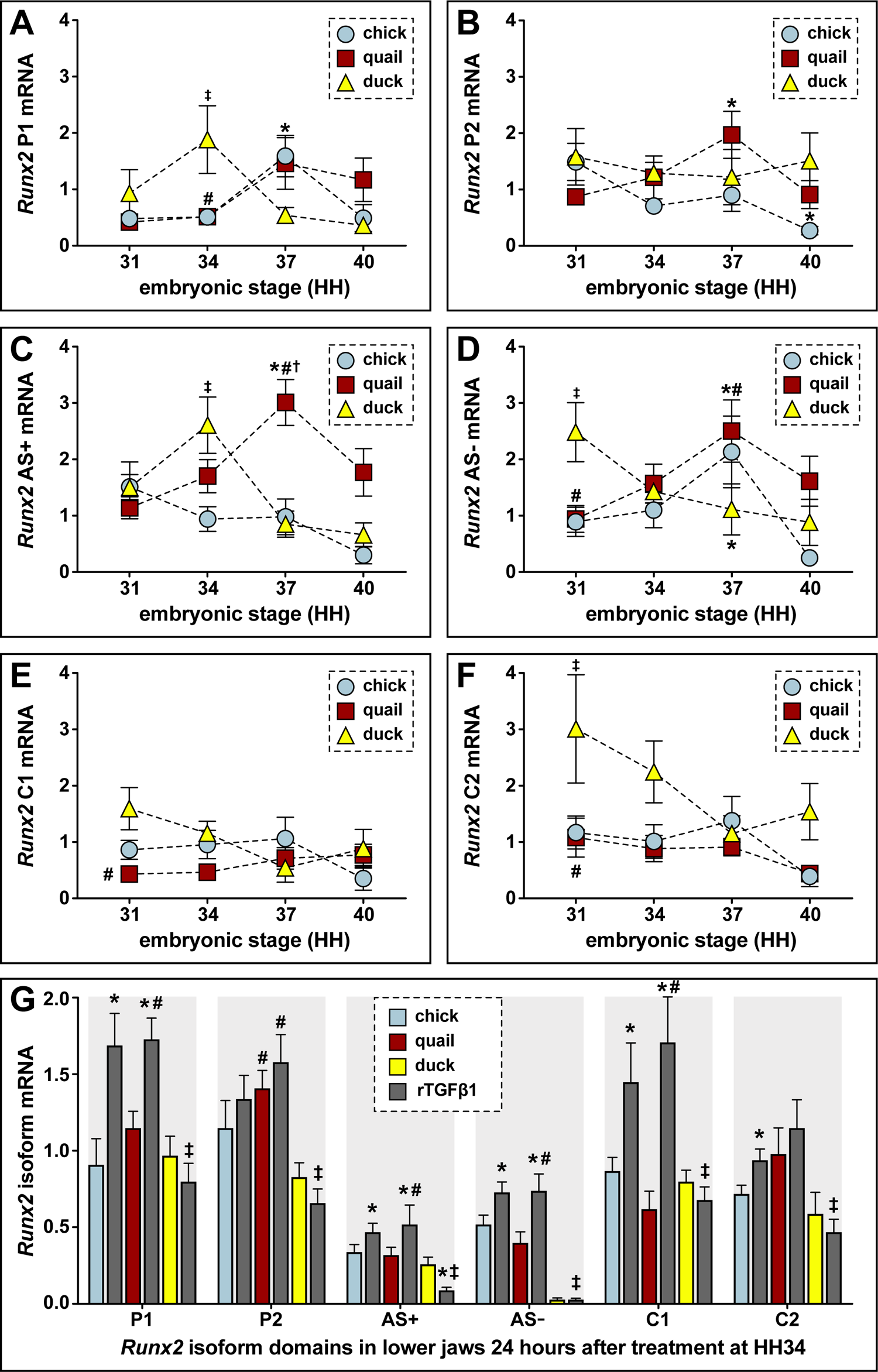
Expression of *Runx2* isoforms containing different functional domains. Relative mRNA **e**xpression levels of all isoforms utilizing P1 (i.e., 1A1, 1A2, 1B1, 1B2), P2 (i.e., 2A1, 2A2, 2B1, 2B2), AS+ (i.e., 1A1, 1A2, 2A1, 2A2), AS-(i.e., 1B1, 1B2, 2B1, 2B2), C1 (i.e., 1A1, 1B1, 2A1, 2B1), and C2 (i.e., 1A2, 1B2, 2A2, 2B2) as measured by qPCR in the jaw primordia of chick, quail, and duck, at HH31, HH34, HH37, and HH40. **A)** P1 increases 3.3-fold in chick (blue, n = 6) and 3.5-fold in quail (red square, n = 9) from HH31 to HH37; chick decreases at HH40. P1 does not change from HH31 to HH34 in duck (yellow triangle, n = 9) but is higher compared to quail at HH34. **B)** P2 decreases 5.5-fold in chick at HH40 and increases 2-fold in quail at HH37. Duck levels do not change over time and no differences are observed among species. **C)** AS+ remains unchanged but decreases in chick by HH40. AS+ increases 3-fold in quail from HH31 to HH37 and is elevated compared to chick and duck at HH37. AS+ decreases 2.2-fold in duck from HH34 to HH40. **D)** AS-increases 2-fold in chick and 2.5-fold in quail at HH37, and quail levels are higher than duck at HH37. AS-decreases 2.8-fold in chick at HH40; and decreases 2.3-fold in duck at HH37, but levels are higher in duck compared to quail and chick at HH31. **E)** C1 does not change over time in any species. **F)** C2 is unchanged in chick and quail over time. C2 decreases 2-fold in duck at HH37 and is higher in duck compared to chick and quail at HH31. * denotes significant difference (p ≤ 0.05) from HH31 within each group, # denotes significant difference (p ≤ 0.05) between quail and duck at same stage, † denotes significant difference (p ≤ 0.05) between chick and quail, ‡ denotes significant difference (p ≤ 0.05) between chick and duck. **G)** Chick (blue), quail (red), and duck (yellow) lower jaws treated in culture with 25 ng/ml rTGFβ1 for 24 hours. rTGFβ1 (gray) induces isoforms utilizing P1 1.9-fold in chick and 1.5-fold quail; AS+ 1.4-fold in chick and 1.6-fold in quail; AS-1.4-fold in chick and 1.8-fold in quail; and C1 1.7-fold in chick and 2.8-fold in quail. rTGFβ1 induces C2 1.3-fold in chick but has no effect in quail or duck. No changes are observed for any domain in duck following treatment with rTGFβ1. Differences between quail and duck in response to rTGFβ1 treatment are observed for P1, P2, AS+, AS-, and C1, but not C2. *denotes significance at p ≤ 0.05 compared to vehicle control, # denotes significance at p ≤ 0.05 between quail and duck at the same stage, † denotes significance at p ≤ 0.05 between chick and quail, ‡ denotes significance at p ≤ 0.05 between chick and duck.

### Runx2 Isoforms are Differentially Expressed in Response to TGFβ Signaling

*Runx2* is a known target of signaling by the TGFβ superfamily (Lee et al., 2000; Alliston et al., 2001; Lee et al., 2002; Ito and Miyazono, 2003; Kang et al., 2005; Derynck et al., 2008). Thus, we tested if TGFβ signaling can differentially regulate expression of individual *Runx2* isoforms, and if this expression is species-specific. We harvested jaw primordia from chick, quail, and duck at HH34 and treated them in culture with rTGFβ1. Twenty-four hours after treatment with rTGFβ1, we observe a 1.9-fold increase in isoforms utilizing the P1 domain in chick (p ≤ 0.001), a 1.5-fold increase for quail (p ≤ 0.02), and no change for duck relative to controls (**Figure 5G**). Moreover, expression of P1 is elevated 2.2-fold in chick (p ≤ 0.0001) and 2.3-fold in quail (p ≤ 0.0001) compared to duck. P2 is not affected by rTGFβ1 for any species, although quail is 2.5-fold higher than duck (p ≤ 0.0001) overall. Isoforms containing the AS+ domain increase 1.4-fold in chick (p ≤ 0.04) and 1.6-fold in quail (p ≤ 0.05), whereas duck decreases 3.6-fold (p ≤ 0.05). Jaw primordia treated with rTGFβ1 showed 23-fold higher AS+ expression in chick (p ≤ 0.0001) and 25-fold higher in quail (p ≤ 0.0001) compared to duck. Isoforms defined by the AS-domain increase 1.4-fold in chick (p ≤ 0.05) and 1.8-fold in quail (p ≤ 0.001); we observed no change in duck. AS-isoforms are 24-fold higher in chick (p ≤ 0.0001) and 24-fold in quail (p ≤ 0.0001) relative to duck. Isoforms containing the C1 domain increase by 1.7-fold in chick (p ≤ 0.05) and 2.8-fold in (p ≤ 0.0002) with no observed change in duck. Following rTGFβ1 treatment, C1 isoforms increase by 2.3-fold in chick (p ≤ 0.005) and 2.8-fold in quail (p ≤ 0.0002) relative to duck. Treatment with rTGFβ1 does not change C2 expression in chick, quail, or duck.

By taking the differential response of each *Runx2* domain into account, we conclude that isoforms 1A1 and 1B1 are regulated by TGFβ signaling in chick and quail since we observe a differential response in the P1, AS+, AS-, and C1 domains. In duck however, we do not observe any TGFβ-mediated effects on 1A1, 2A1, 1A2, or 2A2, and based on the response of isoforms containing the AS+ domain, we find that TGFβ signaling likely has a repressive effect on the expression of 1A1, 2A1, 1A2, 2A2. Such findings are consistent with our prior work demonstrating that chick and quail have elevated TGFβ signaling relative to duck during these same stages of jaw development, that duck is inherently less sensitive to TGFβ signaling than chick or quail, and that TGFβ signaling can repress target genes in duck but activate them in chick and quail (Smith et al., 2020). Additionally, these species-specific differences may also reflect the ability of TGFβ signaling to regulate splicing factors and alternative splicing events that promote the expression of isoforms involved in tissue remodeling and a broad range of normal and abnormal cellular behaviors (Zhao and Young, 1995; Cordray and Satterwhite, 2005; DuRaine et al., 2011; Shirakihara et al., 2011; Hallgren et al., 2012; Tripathi et al., 2016; Abou-Ezzi et al., 2019; Chen et al., 2021).

### Runx2 Isoforms Can Activate or Repress the Expression of Target Genes

To determine the extent to which each isoform of *Runx2* affects the transcription of target genes we performed *in vitro* and *in ovo* overexpression experiments and measured transcriptional activity using qPCR analyses of gene expression and/or luciferase assays. For our *in vitro* experiments, we first tested if chick and duck fibroblast cell lines express *Runx2* isoforms by using primers to amplify all of the isoforms containing the P1, P2, AS+, AS-, C1, and C2 domains. When we measure the total amount of *Runx2* mRNA we find that chick cells express 2-fold higher levels (p ≤ 0.0001) than duck cells (**Figure 6A**), which is consistent with what we observe in the jaw primordia of chick versus duck at HH37 (**Figure 3M**). Similarly, chick cells show 10-fold higher expression (p ≤ 0.0001) of P1 domains, 30-fold higher expression (p ≤ 0.0001) of P2 domains, and 3-fold higher expression (p ≤ 0.0001) of C2 isoforms compared to that observed in duck cells. In both chick and duck cells, we observe more or less equivalent expression levels for isoforms containing the AS+ and C1 domains and find that the AS-isoforms are expressed at low levels. Thus, chick and duck cell lines exhibit species-specific differences in endogenous levels of total *Runx2* and of *Runx2* isoforms.

**Figure 6.**
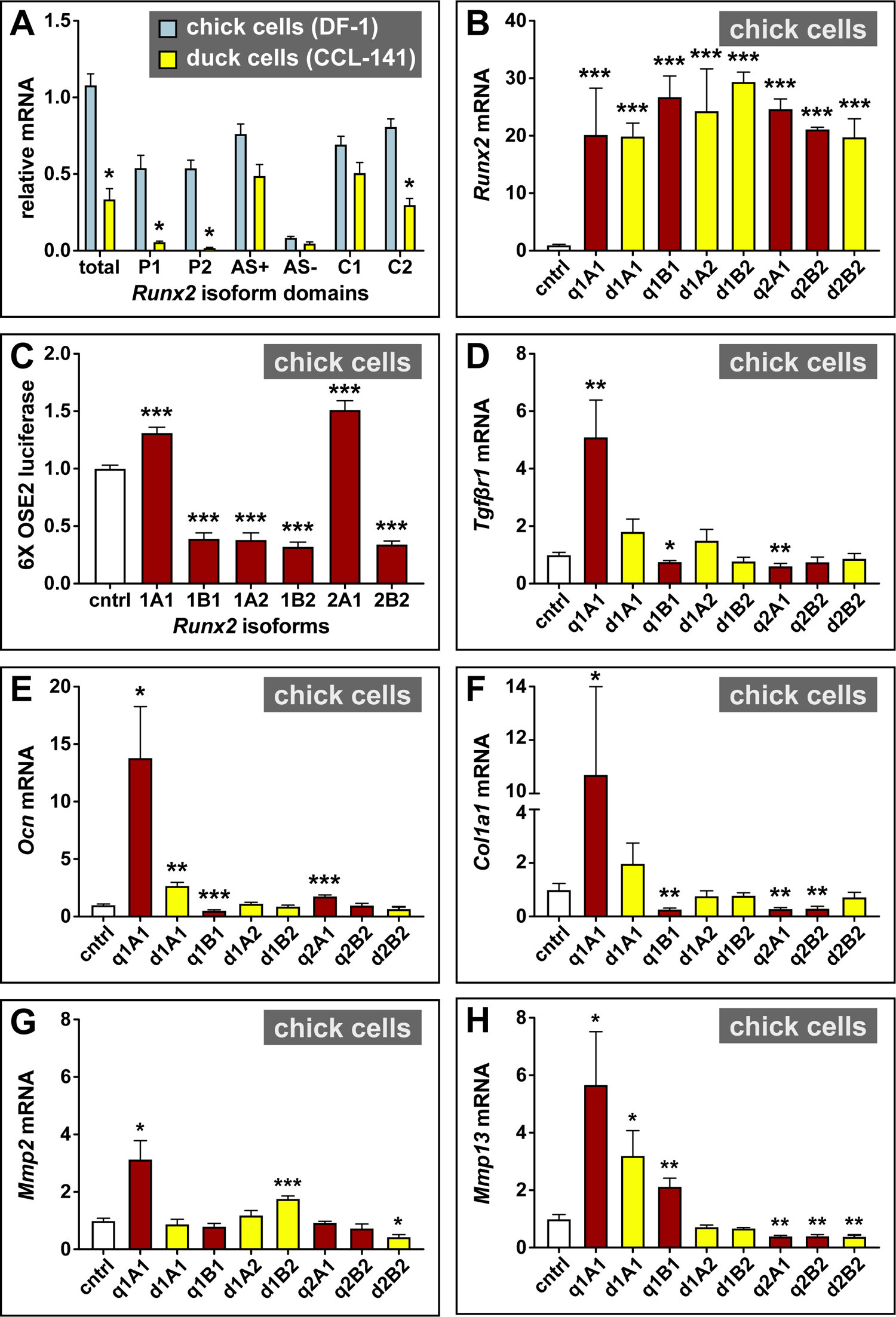
Expression and transcriptional activity of *Runx2* isoforms in cells. **A)** Relative expression of all isoforms utilizing P1 (i.e., 1A1, 1A2, 1B1, 1B2), P2 (i.e., 2A1, 2A2, 2B1, 2B2), AS+ (i.e., 1A1, 1A2, 2A1, 2A2), AS-(i.e., 1B1, 1B2, 2B1, 2B2), C1 (i.e., 1A1, 1B1, 2A1, 2B1), and C2 (i.e., 1A2, 1B2, 2A2, 2B2) as measured by qPCR in chick (blue) and duck (yellow) cells (n = 6 wells per species). Total *Runx2* is 2-fold lower in duck cells compared to chick. P1, P2, and C2 are lower in duck cells compared to chick. AS-is lowest overall compared to other domains. * denotes significance at p ≤ 0.05 for duck cells compared to chick. **B)** Total *Runx2* following overexpression of individual quail (red) and duck (yellow) *Runx2* isoforms in chick cells as measured by qPCR. Each isoform is expressed more than 20-fold higher compared to the empty vector control (white). *** denotes significance at p ≤ 0.001 compared to empty vector control. **C)** Chick cells were co-transfected with an overexpression plasmid containing individual *Runx2* isoforms and a plasmid containing six repeating *Runx2* binding elements (6X OSE2) attached to a luciferase reporter. After 24 hours, *Runx2* isoforms 1A1 and 2A1 induce OSE2 activity by 1.5-fold, whereas 1B1, 1A2, 1B2, and 2B2 decrease activity by 2-fold (n = 18 wells per isoform). **D)** *Tgfβr1* following overexpression of quail and duck *Runx2* isoforms (n = 8 wells per isoform). Quail 1A1 induces *Tgfβr1* by 5-fold compared to empty vector control. Quail 1B1 and 2A1 repress *Tgfβr1* by 2-fold. **E)** Quail 1A1 induces *Ocn* by 15-fold compared to control, while duck 1A1 induces *Ocn* by 3-fold. Quail 2A1 induces *Ocn* by 2-fold, while quail 1B1 decreases *Ocn* by 3-fold. **F)** Quail 1A1 induces *Col1a1* by 10-fold and quail 1B1, 2A1, and 2B2 decrease *Col1a1* by 3-fold. **G)** Quail 1A1 induces *Mmp2* by 2.5-fold and duck 1B2 induces *Mmp2* by 2-fold. Duck 2B2 represses *Mmp2* by 2-fold. **H)** Quail 1A1 induces *Mmp13* by 6-fold, while duck 1A1 induces *Mmp13* by 3.5-fold. Quail 1B1 induces *Mmp13* by 2-fold. Quail 2A1, 2B2, and duck 2B2 repress *Mmp13* by 2.5-fold. Compared to controls, * denotes significance at p ≤ 0.05, ** denotes significance at p ≤ 0.01, and *** denotes significance at p ≤ 0.001.

To overexpress *Runx2* isoforms in chick and duck cells we utilized a dox-inducible promoter system that we characterized previously (Chu et al., 2020). We confirmed *Runx2* isoform overexpression by mScarlet (RFP) fluorescence and by measuring expression of total *Runx2* mRNA. We find that *Runx2* isoform levels are approximately 20-fold higher (p ≤ 0.0001) following overexpression compared to an empty vector control (**Figure 6B**). To measure the functional effects of different *Runx2* isoforms, we overexpressed 1A1, 1B1, 1A2, 1B2, 2A1, and 2B2 in chick cells that were co-transfected with the 6X OSE2 construct, which contains *Runx2* binding elements attached to a luciferase reporter (Karsenty et al., 1999). Compared to empty vector controls, *Runx2* 1A1 and 2A1 induce OSE2 activity by 1.5-fold (p ≤ 0.0001), while 1B1, 1A2, 1B2, and 2B2 decrease activity by 2-fold (p ≤ 0.0001) (**Figure 6C**). We also measured the functional effects of different quail and duck isoforms on expression of known RUNX2 targets, including *Transforming growth factor β receptor 1* (*Tgfβr1*), *Osteocalcin* (*Ocn*), *Collagen type 1* (*Col1a1*), *Matrix metalloproteinase 2* (*Mmp2*), and *Mmp13*. We find that overexpression of quail 1A1 increases *Tgfβr1* by 5-fold (p ≤ 0.01), while duck 1A1 has no effect on *Tgfβr1*; whereas we observe a 2-fold decrease in *Tgfβr1* for 1B1 (p ≤ 0.02) and 2A1 (p ≤ 0.01) (**Figure 6D**). Overexpression of quail 1A1 increases *Ocn* by 15-fold (p ≤ 0.03), while duck 1A1 increases *Ocn* by 3-fold (p ≤ 0.002); quail 2A1 increases *Ocn* by 2-fold (p ≤ 0.0001); and quail 1B1 decreases *Ocn* by 3-fold (p ≤ 0.0001) (**Figure 6E**). Quail 1A1 increases *Col1a1* by 10-fold (p ≤ 0.04) and duck 1A1 has no significant effect on *Col1a1*, whereas quail 1B1, 2A1, and 2B2 decrease *Col1a1* by 3-fold (p ≤ 0.0001) (**Figure 6F**). Quail 1A1 increases *Mmp2* by 2.5-fold (p ≤ 0.05) and duck 1B2 increases *Mmp2* by 2-fold (p ≤ 0.0001); whereas duck 2B2 decreases *Mmp2* by 2-fold (p ≤ 0.02) (**Figure 6G**). Quail 1A1 increases *Mmp13* by 6-fold (p ≤ 0.02), duck 1A1 increases *Mmp13* by 3.5-fold (p ≤ 0.05), and quail 2B1 increases *Mmp13* by 2-fold (p ≤ 0.002); whereas quail 2A1 decreases *Mmp13* by 2.5-fold (p ≤ 0.002), quail 2B2 decreases *Mmp13* by 2.5-fold (p ≤ 0.002), and duck 2B2 decreases *Mmp13* by 2.5-fold (p ≤ 0.001) (**Figure 6H**).

Overall, our *in vitro* isoform overexpression experiments demonstrate that in general 1A1 is the most transcriptionally activating *Runx2* isoform while 1B1 and 2B2 are the most repressive based on the response of the OSE2 binding element and the differential expression of target genes. However, the effects on some genes do vary such as in the case of 2A1 activating OSE2 but repressing everything else except *Mmp2*; 1B2 activating *Mmp2*; or 1B1 activating *Mmp13* but repressing OSE2, *Tgfβr1*, *Ocn*, and *Col1a1*. This gene-specific variation indicates that context is important and that other types of coincident regulation may be contributing to the effects of different isoforms.

### RUNX2 Isoforms Differentially Regulate Target Genes During Jaw Development

To understand the regulatory effects of different functional domains of *Runx2* within the context of jaw development, we overexpressed the 1A1 and 2B2 isoforms in quail and duck embryos. We focused on these two isoforms because they are the most distinct from one another with each defined by either an alternative promoter (P1 or P2), the presence or absence of the domain encoded by exon 5 (AS+ or AS-), or an alternative C-terminus (C1 or C2) (**Figure 4A**). To restrict overexpression to late stages of jaw development, we utilized a tool we previously developed that combines a piggyBac transposon system, which integrates into the host genome and expresses GFP, and a dox-inducible promoter, which drives overexpression of RFP and a gene of interest (Chu et al., 2020). We unilaterally electroporated presumptive NCM at HH8.5 so that the non-electroporated contralateral side could serve as an internal control (**Figure 7A**), treated embryos with dox at HH34 (**Figure 7B**), and then examined the extent of isoform overexpression at HH37 by screening embryos for RFP using epifluorescent or confocal microscopy (**Figure 7C**). Unilateral overexpression of total *Runx2* mRNA was confirmed by qPCR. We observe a 4.5-fold (p ≤ 0.01) increase of 1A1 in quail and a 4.5-fold (p ≤ 0.02) increase in duck, and a 4.4-fold (p ≤ 0.01) increase of 2B2 in quail and a 7.3-fold (p ≤ 0.003) increase in duck (**Figure 7D**).

**Figure 7.**
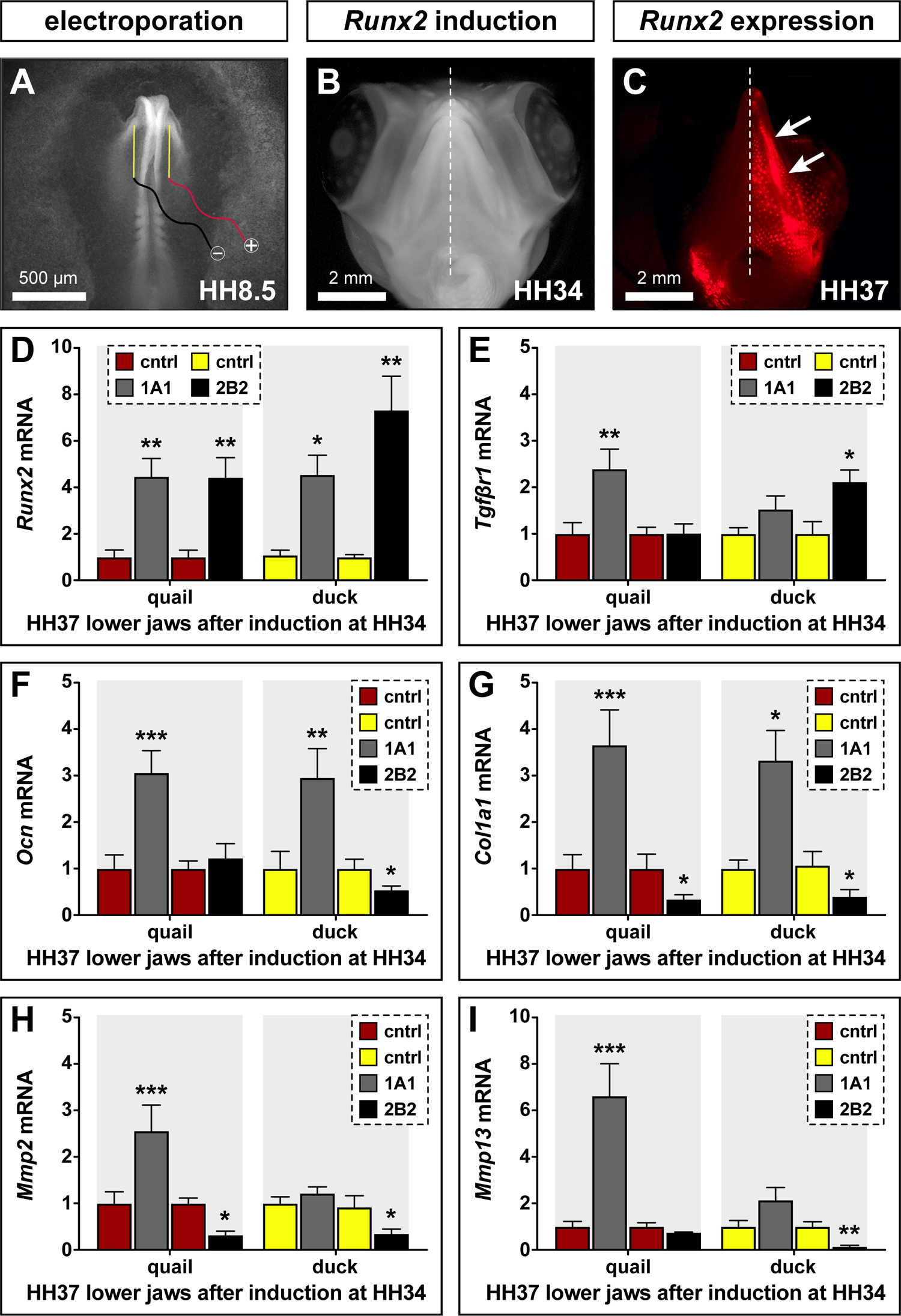
Transcriptional activity of *Runx2* isoforms 1A1 versus 2B2 *in ovo*. **A)** Presumptive NCM was electroporated unilaterally in quail and duck embryos as HH8.5 with species-matched 1A1 or 2B2 overexpression constructs. **B)** At HH34, embryos were treated with dox for 48 hours. **C)** At HH37, specimens were screened by epifluorescence for positive unilateral RFP signal (arrows). **D)** Total *Runx2* following overexpression of 1A1 (gray) and 2B2 (black) in quail (red) and duck (yellow) lower jaws as measured by qPCR. *Runx2* is 4.5-fold higher in quail and duck following 1A1 overexpression compared to the contralateral side; and 4.5-fold higher in quail and 7.3-fold higher in duck following 2B2 overexpression. **E)** *Tgfβr1* is 2.4-fold higher in quail following 1A1 overexpression, and 2.1-fold higher in duck following 2B2 overexpression. **F)** *Ocn* is 3-fold higher in quail and duck following 1A1 overexpression, and 1.9-fold lower in duck following 2B2 overexpression. **G)** *Col1a1* is 3.6-fold higher in quail and 3.3-fold in duck following 1A1 overexpression, and 2.9-fold lower in quail and 2.5-fold lower in duck following 2B2 overexpression. **H)** *Mmp2* is 2.6-fold higher in quail following 1A1 overexpression, and 3.1-fold lower in quail and 2.9-fold lower in duck following 2B2 overexpression. **I)** *Mmp13* is 6.6-fold higher in quail following 1A1 overexpression and 7.1-fold lower in duck following 2B2 overexpression. Compared to contralateral control side, * denotes significance at p ≤ 0.05, ** denotes significance at p ≤ 0.01, and *** denotes significance at p ≤ 0.001. n = 7 to 11 for each treatment group.

With regard to the regulation of RUNX2 target genes and relative to contralateral controls, 1A1 overexpression in quail increases *Tgfβr1* by 2.4-fold (p ≤ 0.006) but does not affect *Tgfβr1* in duck (**Figure 7E**). 2B2 overexpression does not affect *Tgfβr1* in quail but increases *Tgfβr1* in duck by 2.1-fold (p ≤ 0.04). 1A1 overexpression increases *Ocn* by 3-fold in both quail (p ≤ 0.001) and duck (p ≤ 0.002) whereas 2B2 overexpression does not affect *Ocn* in quail but decreases *Ocn* by 1.9-fold (p ≤ 0.05) in duck (**Figure 7F**). 1A1 overexpression increases *Col1a1* by 3.6-fold in quail (p ≤ 0.0001) and 3.3-fold in duck (p ≤ 0.03) whereas 2B2 overexpression decreases *Col1a1* by 2.9-fold in quail (p ≤ 0.03) and 2.5-fold in duck (p ≤ 0.05) (**Figure 7G**). 1A1 overexpression increases *Mmp2* by 2.6-fold in quail (p ≤ 0.0001) but does not affect *Mmp2* in duck, whereas 2B2 overexpression decreases *Mmp2* by 3.1-fold in quail (p ≤ 0.04) and 2.9-fold in duck (p ≤ 0.04) (**Figure 7H**). 1A1 increases *Mmp13* by 6.6-fold in quail (p ≤ 0.0001) but does not change *Mmp13* in duck whereas 2B2 overexpression does not alter *Mmp13* in quail but decreases *Mmp13* by 7.1-fold in duck (p ≤ 0.002) (**Figure 7I**). In sum, we find that the 1A1 isoform transcriptionally activates all five target genes examined in quail (i.e., *Tgfβr1*, *Ocn*, *Col1a1*, *Mmp2*, and *Mmp13*) but only the two osteogenic genes in duck (i.e., *Ocn* and *Col1a1*). The 2B2 isoform represses two genes in quail (i.e., *Col1a1* and *Mmp2*) whereas 2B2 represses four genes in duck (i.e., *Ocn*, *Col1a1*, *Mmp2*, and *Mmp13*) and activates one (i.e, *Tgfβr1*). Such species-specific differences in the ways 1A1 versus 2B2 feed into the regulatory landscape suggest that developmental context helps determine the response of target genes to RUNX2 isoforms, that the alternatively spliced domains of RUNX2 (i.e., P1, P2, AS+, AS-, C1, and C2) have distinct functional effects, and that interactions at the level of promoters and enhancers may play an essential role in *Runx2*-mediated programs for bone deposition and resorption.

### Runx2 Isoforms Differentially Regulate the Mmp13 Promoter

As a proof-of-principle to understand mechanisms through which RUNX2 isoforms interact with target genes, we focused on the *Mmp13* promoter. The *Mmp13* promoter in particular, is well characterized and known to be regulated by RUNX2 (Pendas et al., 1997; Wang et al., 2004; Chen et al., 2012a; Meyer et al., 2016; Takahashi et al., 2017; Young et al., 2019; Smith et al., 2020). Additionally, results from both our *in vitro* and *in ovo* experiments demonstrate that *Mmp13* is among the most differentially regulated genes of the ones we examined (in terms of fold change) and shows significant and dramatic species-specific differences in its response to the same isoforms. For example, we find that 1A1 can activate *Mmp13* in quail but not duck, or 2B2 can repress *Mmp13* in duck but not quail (**Figure 7I**). In a prior study, we cloned and characterized the first −2 kb of the *Mmp13* promoters of chick, quail, and duck and identified all of the RUNX2 binding elements (Smith et al., 2020). While three RUNX2 binding elements are present in the *Mmp13* promoters of all three species, they are found in different locations except for one common site in the most proximal region (**Figure 8A**).

**Figure 8.**
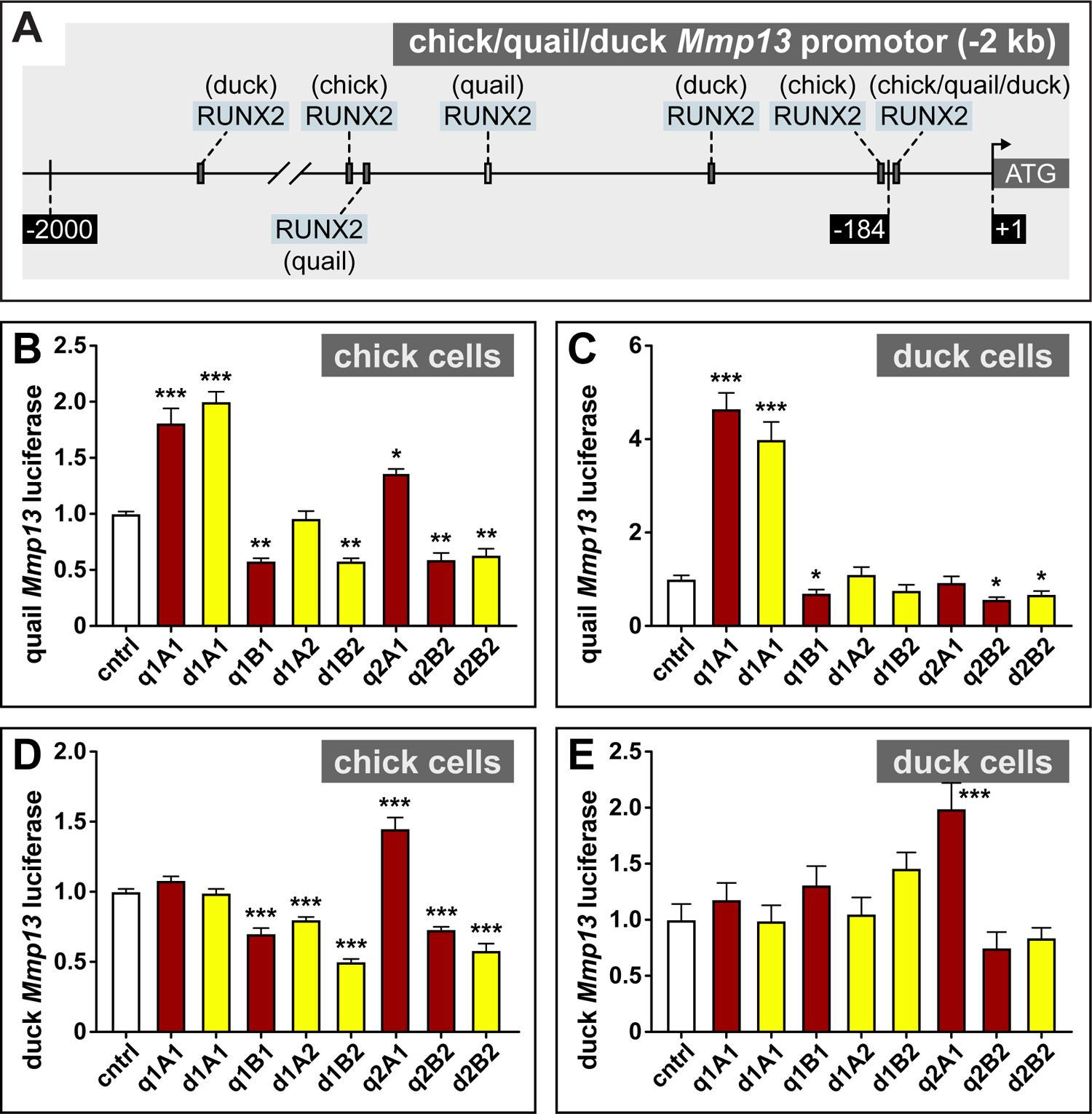
Regulation of the *Mmp13* promoter by *Runx2* isoforms. **A)** Schematic of a −2 kb region of the chick, quail, and duck *Mmp13* promoter upstream from the transcriptional start site (ATG). Chick, quail, and duck each have two RUNX2 binding elements in different locations and share a common site in the proximal promoter. **B)** Chick cells were co-transfected with an overexpression plasmid containing individual quail (red) and duck (yellow) isoforms and a plasmid containing −2 kb of the *Mmp13* promoter attached to luciferase. After 24 hours, quail and duck 1A1 induce *Mmp13* promoter activity by 2-fold; quail 2A1 induces *Mmp13* promoter activity by 1.5-fold; and quail 1B1, duck 1B2, and quail and duck 2B2 repress *Mmp13* promoter activity by 2-fold relative to empty vector controls (white). **C)** In duck cells, quail and duck 1A1 induce quail *Mmp13* promoter activity by 4-fold; and quail 1B1 and quail and duck 2B2 repress *Mmp13* promoter activity by 2-fold. **D)** In chick cells, quail 2A1 induces duck *Mmp13* promoter activity by 1.5-fold; and quail 1B1, duck 1A2, duck 1B2, and quail and duck 2B2 repress *Mmp13* promoter activity by 1.4-fold. **E)** In duck cells, quail 2A1 induces duck *Mmp13* promoter activity by 2-fold. Compared to control, * denotes significance at p ≤ 0.05, ** denotes significance at p ≤ 0.01, and *** denotes significance at p ≤ 0.001. n = 18 to 36 for each treatment group.

We cloned −2 kb fragments of the quail and duck *Mmp13* promoters into luciferase expression constructs and co-transfected these and different RUNX2 isoform overexpression constructs into chick and duck cells. We find that in chick cells (**Figure 8B**), overexpressing quail 1A1 and duck 1A1 induces quail *Mmp13* promoter activity by 2-fold (p ≤ 0.0001), and overexpressing quail 2A1 induces quail *Mmp13* promoter activity by 1.5-fold (p ≤ 0.01). In contrast, overexpressing quail 1B1 (p ≤ 0.002), quail 1B2 (p ≤ 0.002), quail 2B2 (p ≤ 0.003), and duck 2B2 (p ≤ 0.01) represses quail *Mmp13* promoter activity by 2-fold. In duck cells, we observe a similar pattern but with substantially higher levels of induction (**Figure 8C**). Overexpressing quail 1A1 and duck 1A1 induces quail *Mmp13* promoter activity by 4-fold (p ≤ 0.0001). In contrast, overexpressing quail 1B1 (p ≤ 0.05), quail 2B2 (p ≤ 0.05), and duck 2B2 (p ≤ 0.05) represses quail *Mmp13* promoter activity by 2-fold. In chick cells, overexpressing quail 1A1 and duck 1A1 has no effect on duck *Mmp13* promoter activity whereas overexpressing quail 2A1 induces duck *Mmp13* promoter activity by 1.5-fold (p ≤ 0.0001) (**Figure 8D**). In contrast, overexpressing quail 1B1 (p ≤ 0.02), duck 1A2 (p ≤ 0.0001), duck 1B2 (p ≤ 0.0001), quail 2B2 (p ≤ 0.0004), and duck 2B2 (p ≤ 0.0001) represses duck *Mmp13* promoter activity by 1.4-fold. In duck cells, overexpressing 2A1 induces duck *Mmp13* promoter activity by 2-fold (p ≤ 0.0002) (**Figure 8E**). Overall, we find certain RUNX2 isoforms activate the *Mmp13* promoter while others repress activity, but developmental context and species-specific promoter elements also appear to have an effect.

### SNPs by a RUNX2 Binding Element Affect Isoform-Mediated Activity of Mmp13

To determine the extent to which the RUNX2 binding elements account for the differential and species-specific response of *Mmp13* to RUNX2 isoform overexpression, we attached luciferase to a −184 bp fragment of the proximal *Mmp13* promoter containing only one RUNX2 binding element in chick and quail, and a −181 bp fragment containing only one RUNX2 binding element in duck. In a prior study we identified two single nucleotide polymorphisms (SNPs) that distinguish chick and quail (i.e., “AG”) from duck (i.e., “CA”) located −160 bp upstream of the *Mmp13* translational start site and immediately adjacent to a RUNX2 binding element within the −184/-181 bp fragments (Smith et al., 2020) (**Figure 9A**). To test if these SNPs affect the differential and species-specific response of the *Mmp13* promoter to various RUNX2 isoforms, we also generated constructs in which we incorporated the chick/quail “AG” SNPs into the duck promoter and the duck “CA” SNPs into the chick/quail promoter.

**Figure 9.**
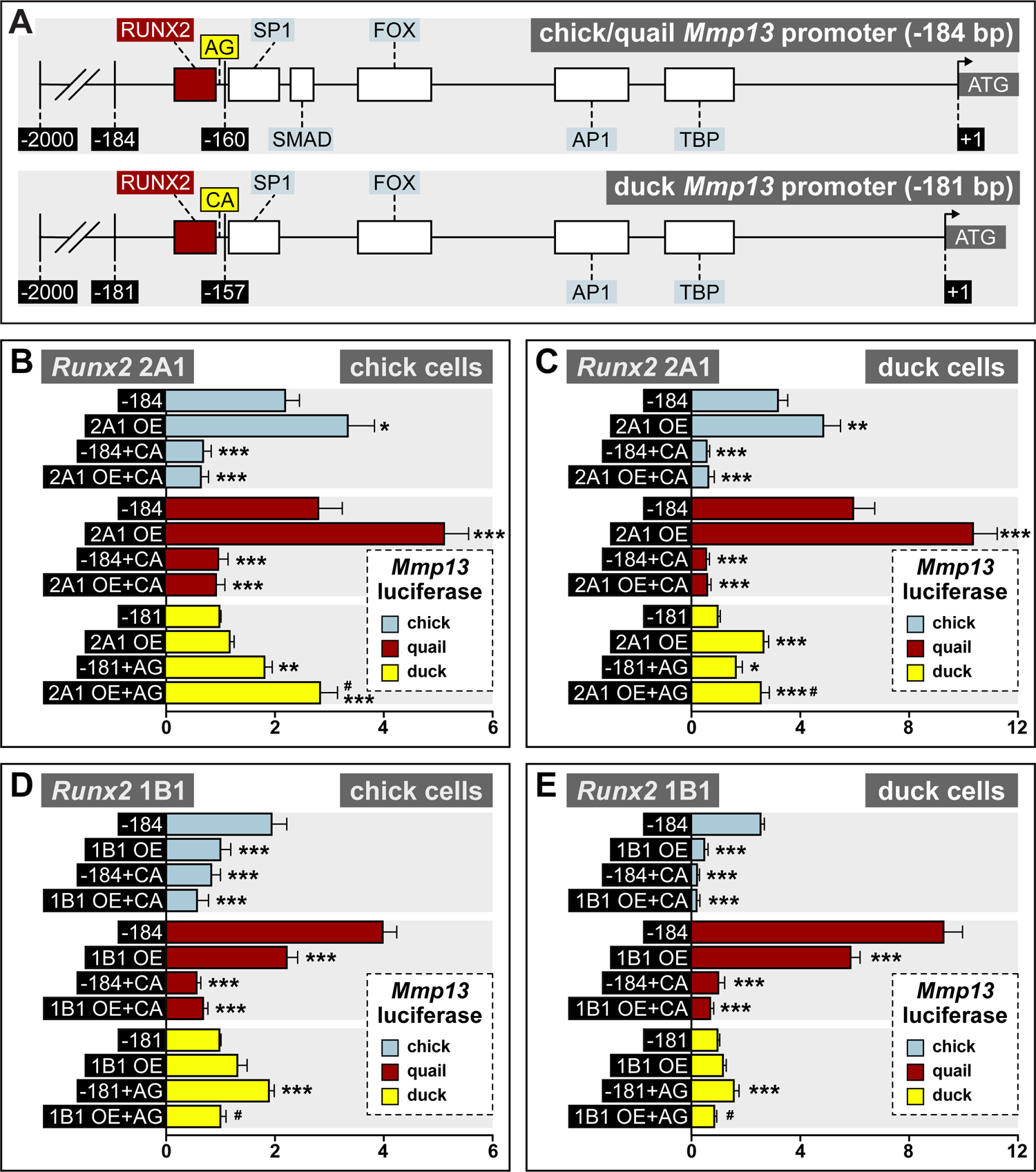
Effects of SNPs on *Mmp13* promoter regulation by *Runx2* isoforms 2A1 and 1B1. **A)** Schematic of the proximal region of the chick and quail (−184 bp), and duck (−181 bp) *Mmp13* promoters. Chick, quail, and duck have equivalent binding elements for TATA Binding Protein (TBP), Activator Protein-1 (AP1), Forkhead Box (FOX), Specificity Protein-1 (SP1), and RUNX2. However, chick and quail contain two SNPs (“AG”) adjacent to the RUNX2 binding element that differ from the duck (“CA”). **B)** Chick cells transfected with chick (blue) or quail (red) *Mmp13* promoters containing either a −184 bp control fragment including the chick/quail “AG” SNPs; a −184 bp fragment including the chick/quail “AG” SNPs plus a *Runx2* 2A1 over-expression (OE) construct; a −184 bp fragment including the duck “CA” SNPs; or a −184 bp fragment including the duck “CA” SNPs plus a 2A1 OE construct. Chick cells were also transfected with the duck (yellow) *Mmp13* promoter containing either a −181 bp control fragment including the duck “CA” SNPs; a −181 bp fragment including the duck “CA” SNPs plus a 2A1 OE construct; a −181 bp fragment including the chick/quail “AG” SNPs; or a −181 bp fragment including the chick/quail “AG” SNPs plus a 2A1 OE construct. In chick cells, 2A1 OE increases activity of the chick *Mmp13* promoter by 1.5-fold and quail promoter by 2.8-fold. Switching duck “CA” SNPs into chick and quail promoters decreases activity by 1.9-fold and by 1.3-fold with 2A1 OE. Switching chick/quail SNPs into the duck promoter leads to an increase in activity by 1.6-fold with 2A1 OE. **C)** In duck cells, 2A1 OE increases activity of the chick *Mmp13* promoter by 1.5-fold and quail promoter by 1.7-fold. Switching the duck SNPs decreases activity in the chick promoter by 4.9-fold and quail by 9.7-fold even with 2A1 OE. 2A1 OE increases activity of the duck promoter by 2.7-fold and switching the “CA” SNPs to the duck promoter increases activity by 1.7-fold and 2.6-fold with 2A1 OE. **D)** In chick cells, 1B1 OE decreases activity of the chick and quail *Mmp13* promoter by 2.0-fold. Switching duck SNPs into the chick promoter decreases activity by 2.3-fold and by 6.9-fold in the quail promoter. Switching chick/quail SNPs into the duck promoter leads to an increase in activity by 1.9-fold but when combined with 1B1 OE, duck promoter activity decreases. **E)** In duck cells, 1B1 OE decreases activity of the chick *Mmp13* promoter by 5-fold and quail promoter by 1.6-fold. Switching duck SNPs into the chick promoter decreases activity by 10.3-fold and by 9.3-fold in the quail promoter and also with 1B1 OE. Switching chick/quail SNPs into the duck promoter increases activity by 1.6-fold but 1B1 OE decreases activity by 1.3-fold. Compared to the endogenous promoter within each species, * denotes significance at p ≤ 0.05, ** denotes significance at p ≤ 0.01, and *** denotes significance at p ≤ 0.001. # denotes significance at p ≤ 0.05 compared to control SNP switch. n = 18 to 36 for each treatment group.

We find that in chick cells and relative to control −184 bp fragments alone, overexpressing 2A1 induces chick *Mmp13* promoter activity by 1.5-fold (p ≤ 0.02) and quail *Mmp13* promoter activity by 2.8-fold (p ≤ 0.0001) (**Figure 9B**). Overexpressing 2A1 has no measurable effect on the duck *Mmp13* promoter. In contrast, switching duck “CA” SNPs into the chick and quail promoter represses the activation observed with the control *Mmp13* promoter by 1.5-fold (p ≤ 0.0001) but this repression is not affected by 2A1 overexpression. Switching chick and quail “AG” SNPs into the duck *Mmp13* promoter increases activity by 1.5-fold (p ≤ 0.003) relative to the control duck −181 bp fragment alone, and overexpressing 2A1 induces activity by 1.9-fold (p ≤ 0.0001), which is similar to what is observed for the control chick and quail *Mmp13* promoters, and which is 1.3-fold (p ≤ 0.0001) higher than the control duck promoter with the chick and quail “AG” SNPs. We observe equivalent trends in duck cells although 2A1 overexpression activates the duck *Mmp13* promoter by 2.7-fold (p ≤ 0.0001) and by 1.7-fold (p ≤ 0.004) when including the chick and quail “AG” SNPs (**Figure 9C**). We find that in chick cells and relative to control −184 bp fragments alone, overexpressing 1B1 reduces both chick and quail *Mmp13* promoter activity by 2.0-fold (p ≤ 0.0001) (**Figure 9D**). Switching duck “CA” SNPs into the chick *Mmp13* promoter reduces activity by 2.3-fold (p ≤ 0.0001) both without and with 1B1 overexpression. Similarly switching duck “CA” SNPs into the quail *Mmp13* promoter reduces activity by 6.9-fold (p ≤ 0.0001) both without and with 1B1 overexpression. In contrast, switching chick and quail “AG” SNPs into the duck *Mmp13* promoter increases activity by 1.9-fold (p ≤ 0.0001) relative to the control duck −181 bp fragment alone, but overexpressing 1B1 reduces expression by 1.9-fold (p ≤ 0.0001) relative to the control duck −181 bp fragment with the “AG” SNPs. We observe equivalent trends in duck cells (**Figure 9E**). We also performed similar experiments with 1B2 and 2B2 and find that overexpressing both isoforms represses chick and quail *Mmp13* promoter activity but not duck *Mmp13* promoter activity. We do observe a change in activity in the duck promoter when 2B2 is overexpressed and the SNPs are switched (**Supplemental Figure 1**).

Thus, even though chick, quail, and duck *Mmp13* promoters contain the same total number of RUNX2 binding elements, species-specific SNPs adjacent to the most proximal RUNX2 binding element appear to be important for interactions with RUNX2 isoforms. Switching chick and quail “AG” SNPs into the duck *Mmp13* promoter upregulates activity, whereas switching the duck “CA” SNPs decreases chick or quail promoter activity, and these effects can be mitigated or enhanced depending on which *Runx2* isoform is overexpressed. For instance, 2A1 normally activates the chick and quail *Mmp13* promoter but not duck. Overexpressing 2A1 with a duck promoter that contains the chick and quail “AG” SNPs, enables 2A1 to become an activating isoform. Conversely, 1B1 normally represses the chick and quail *Mmp13* promoter but activates the duck *Mmp13* promoter. Overexpressing 1B1 with a duck promoter that contains the chick and quail “AG” SNPs, enables 1B1 to become a repressive isoform. Such results suggest that species-specific regulation of *Mmp13* depends upon the way different RUNX2 isoforms interact with the RUNX2 binding element and adjacent SNPs.

### Runx2 Contains Multiple Domains that Direct its Transcriptional Activity

Although the overall structure of *Runx2* has remained highly conserved across vertebrates, and especially within amniotes, our study suggests that the stage- and species-specific deployment of *Runx2* isoforms by NCM regulates the transcriptional activity of target genes in ways that likely contribute to the evolution of the jaw skeleton. Presumably this is because each isoform is composed of distinct functional domains that allow RUNX2 protein to be transcriptionally activating and/or repressive depending on the nature of interactions with target genes. In addition, we find that species-specific polymorphisms in the *Mmp13* promoter also influence the regulatory effects of various isoforms and we expect that this would be the same for many other target genes. We imagine that these regulatory effects can be modulated even further by altering the context such as where, when, and how RUNX2 protein becomes localized; the cell state or cell type in which RUNX2 protein is expressed; and the ways RUNX2 protein becomes modified post-translationally (Schneider, 2018a). Ultimately, much of this context can be defined by the inclusion or exclusion of relevant functional elements in each *Runx2* isoform such as P1 or P2, a nuclear localization signal (NLS), nuclear matrix-targeting signal (NMTS), and various pentapeptide motifs (**Figure 1A**).

For example, expression from the P1 promoter is associated with mesenchymal cells, pre-osteoblasts, chondrocyte precursors, and endochondral ossification while expression from P2 is associated with osteoblasts, hypertrophic chondrocytes, intramembranous ossification, and non-osteogenic expression (Park et al., 2001; Choi et al., 2002; Xiao et al., 2004; Li and Xiao, 2007; Jeong et al., 2008; Okura et al., 2014). In our study, we observe species-specific differences in expression of P1 (**Figure 5A**) but not P2 (**Figure 5B**) among chick, quail, and duck. Perhaps this is related to the fact that the lower jaw of birds ossifies exclusively through intramembranous ossification except for the proximal most region of Meckel’s cartilage, which undergoes endochondral ossification but only at stages later than what we examined (Starck, 1989; Helms and Schneider, 2003; Eames et al., 2004; Mitgutsch et al., 2011; Ealba et al., 2015; Anthwal et al., 2017; Svandova et al., 2020). Other studies on activation and repression by isoforms utilizing P1 versus P2 indicate that both have similar transcriptional potentials (Banerjee et al., 2001). Our results do not support these conclusions since we find the 1A1 isoform activates *Tgfβr1*, *Ocn*, *Mmp2*, and *Mmp13* mRNA expression more than 2A1 (**Figure 6D to 6H**). However, we find that 2A1 is a transcriptional activator of the OSE2 binding site and of the *Mmp13* promoter (**Figure 6C and 6E, Figure 8B to 8E**, **Figure 9B to 9C**). In contrast, another study has found that isoforms utilizing P2 can activate the *Ocn* promoter more than P1, but only when using a higher dose of plasmid for overexpression (Xiao et al., 1999). Isoforms utilizing P1 have also been found to mediate the endothelial-mesenchymal transition (EMT) within the developing heart, which is regulated by TGFβ2 (Tavares et al., 2018), and P2 appears to be associated with the development and differentiation of the tooth germ, whereas P1 is not (Kobayashi et al., 2006). We find that P1 but not P2 is induced by TGFβ1. In a prior study we observed that TGFβ signaling is more active in the lower jaws of quail than duck during later stages of development (Smith et al., 2020), which supports our observations here that quail show higher 1A1 isoform expression than duck (**Figure 4C**). Similarly, other experiments have found that CCAAT/Enhancer Binding-Protein β (C/EBPβ) and Specificity Protein 1 (SP1) upregulate P1 expression (Zhang et al., 2009; Henriquez et al., 2011); less is known about P2 regulation.

Additional elements critical for RUNX2 function include a classic NLS (**Figure 1A**), which was discovered based on its homology to the C-Myc NLS and which partially overlaps alternatively spliced exon 5 and the RHD at its C-terminus (Thirunavukkarasu et al., 1998). A second NLS is found at the N-terminus of the RHD, and mutations in either NLS result in disrupted RUNX2 localization and cleidocranial dysplasia (CCD) (Kim et al., 2006; Hansen et al., 2011). Our results also indicate that the NLS encoded by exon 5 may play an important role, since isoforms that include exon 5 appear to be more transcriptional activating (1A1, 2A1) than isoforms with exon 5 spliced out (1B1, 2B2) (**Figure 6** and **Figure 9**). Another essential element is the NMTS located in C1, which forms a loop-turn-loop structure (Zeng et al., 1997; Tang et al., 1999) that interacts directly with SMADS and directs them to the nuclear matrix (Zaidi et al., 2002). The NMTS domain is required for proper nuclear localization of RUNX2, which normally appears punctate but becomes diffuse following a loss of the NMTS (Zaidi et al., 2001; Harrington et al., 2002). Moreover, disruptions to the structure of the NMTS reduce RUNX2 activity and inhibit osteogenic differentiation (Javed et al., 2005; Zaidi et al., 2006), and missense mutations in the NMTS cause CCD (Kim et al., 2006). Finally, C1 also contains two pentapeptide motifs, GASEL and VWRPY. The GASEL motif is found at the start of exon 8 and the VWRPY motif is found at the end followed directly by a stop codon. The GASEL domain is sufficient to induce transcriptional activation whereas the VWRPY motif interacts with transcriptional co-repressors (Aronson et al., 1997; Walrad et al., 2010; Kim et al., 2020). In contrast, C2 remains poorly characterized (Terry et al., 2004) and we find C2 to be highly variable across the species we examined (**Supplemental Table 4**).

### Alternative Splicing Serves as a Key Evolutionary Developmental Mechanism

Alternative splicing is an evolutionarily conserved process yielding mRNA isoforms that can encode a diversity of protein variants with distinct activity, stability, localization, and function (Black, 2003; Matlin et al., 2005). Alternative splicing events are often differentially regulated across tissues and during development, as well as among individuals and populations, suggesting that individual isoforms may play distinct spatial or temporal roles. Estimates for human tissues indicate that up to 92-95% of multi-exon genes express multiple isoforms (Pan et al., 2008; Wang et al., 2008). Variation in isoforms appears to evolve more rapidly than actual changes in gene expression among vertebrates and therefore provides a powerful mechanism for expanding the regulatory and functional repertoire of genes (Barbosa-Morais et al., 2012; Merkin et al., 2012; Reyes et al., 2013). For example, comparisons between embryonic and adult tissues in the brain have revealed that 31% of genes are differentially regulated by alternative splicing even though their total expression levels remain unchanged (Dillman et al., 2013). Alternative splicing is under precise temporal control (Jessell, 2000; Molyneaux et al., 2007; Raj and Blencowe, 2015; Weyn-Vanhentenryck et al., 2018) and can play an essential role in determining how tissues acquire their identity during cell lineage commitment and differentiation (Wang et al., 2008; Salomonis et al., 2010; Gabut et al., 2011; Baralle and Giudice, 2017).

Increased levels and complexity of alternative splicing especially in higher vertebrates can underlie phenotypic change (Kim et al., 2007; Nilsen and Graveley, 2010; Chen et al., 2012c) and drive rapid adaptive radiation such as in the jaws of cichlid fish (Singh et al., 2017). Alternative splicing is critical for normal development of the craniofacial complex and dysregulation of splicing factors can disrupt patterning (Bélanger et al., 2018; Hooper et al., 2020; Lee et al., 2020; Wood et al., 2020). Multiple craniofacial disorders have spliceosomopathies as their primary etiology (Bernier et al., 2012; Lehalle et al., 2015; Merkuri and Fish, 2019). For example, deletion of splicing factors in NCM can cause cleft palate and a disrupted craniofacial skeleton (Cibi et al., 2019; Beauchamp et al., 2020). Such work emphasizes the increasing importance of understanding the role of alternative splicing during embryonic development in the context of both disease and evolution. As a case in point, our study demonstrates how a clinically relevant transcription factor like *Runx2*, which consists of domains that are highly conserved across vertebrates, can diversify its regulatory abilities and functional effects by deploying different combinations of activating and repressive isoforms in time and space throughout development, and by modulating this deployment among species during evolution.

## ACKNOWLEDGEMENTS

We thank T. Alliston and N. Dole for helpful discussions; J. Aggleton, Z. Vavrušová, A. Nguyen, and P. Asfour for technical assistance. We thank T. Dam at AA Lab Eggs. The pmScarlet-i_C1 was a gift from Dorus Gadella (Addgene, #85044). The AAVS1 Puro Tet3G 3xFLAG Twin Strep was a gift from Yannick Doyon (Addgene, # 92099). The pKanCMV-mClover3-mRuby3 was a gift from Michael Lin (Addgene, #74252). The pCAG-Cre-IRES2-GFP was a gift from Anjen Chenn (Addgene, #26646). The pCMV-hyPBase was provided by the Wellcome Trust Sanger Institute. The mNeonGreen was provided by Allele Biotechnology & Pharmaceuticals. This work was supported in part by the UCSF Biological Imaging Development Core (BIDC); the UCSF Core Center for Musculoskeletal Biology and Medicine (CCMBM) through NIAMS P30 AR066262; NIDCR F31 DE027283 to S.S.S.; and NIDCR R01 DE016402 and R01 DE025668, and NIH Office of the Director S10 OD021664 to R.A.S.

## AUTHOR CONTRIBUTIONS

S.S.S, D.B.C., and R.A.S. conceived of the project; S.S.S, D.B.C., and R.A.S. designed the experiments; S.S.S, D.B.C., T. Q., T.H., A.J.L, and G.K. performed the experiments; S.S.S, D.B.C., T.Q., T.H., A.J.L, G.K., and R.A.S. analyzed the data; and S.S.S, D.B.C., and R.A.S. co-wrote the manuscript.

## DATA AVAILABILITY

All datasets and constructs will be made publicly available at the time of publication.

---

**Supplemental Figure 1.**
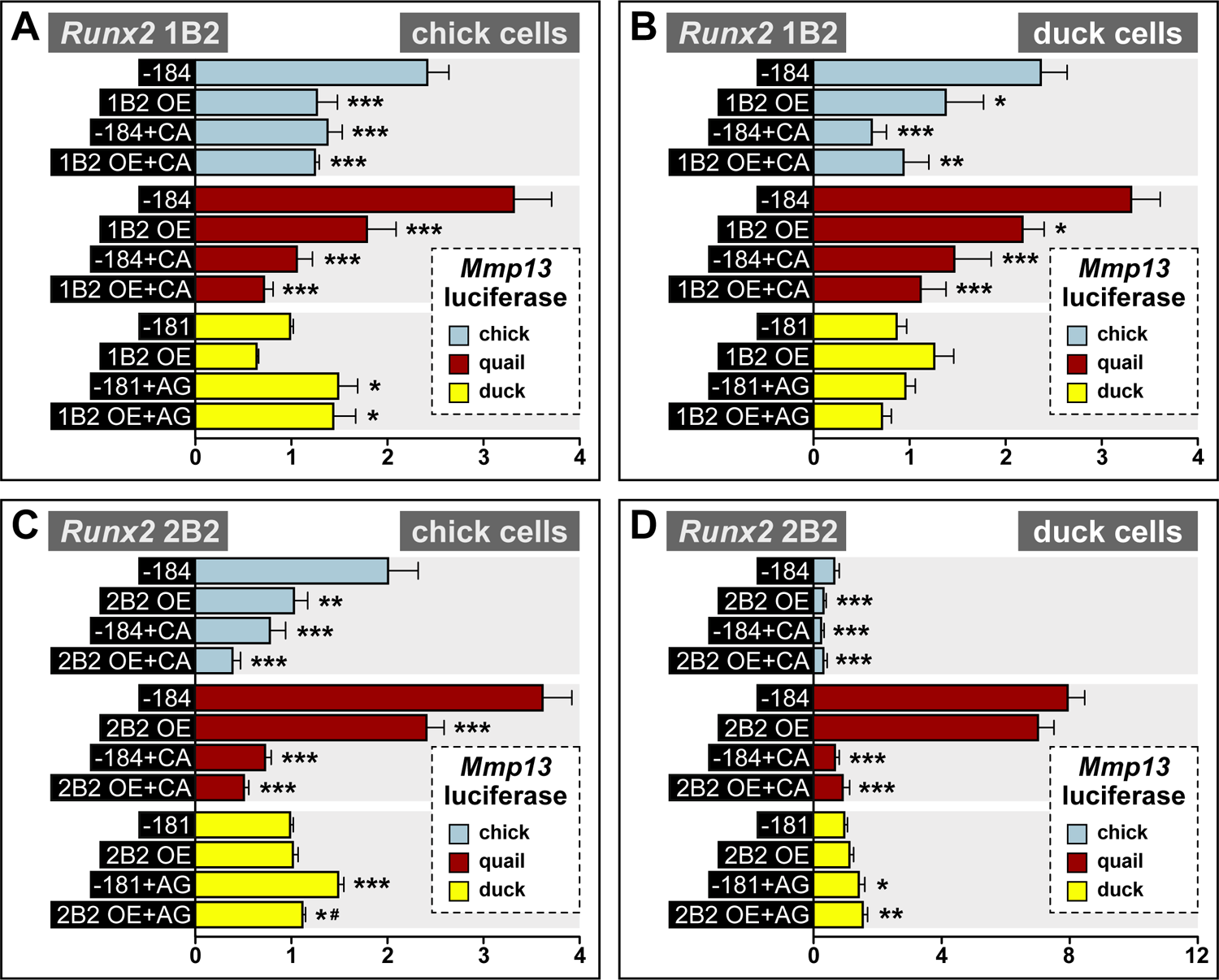
Effects of SNPs on *Mmp13* promoter regulation by *Runx2* isoforms 1B2 and 2B2. **A)** Chick cells transfected with chick (blue) or quail (red) *Mmp13* promoters containing either a −184 bp control fragment including the chick/quail “AG” SNPs; a −184 bp fragment including the chick/quail “AG” SNPs plus a *Runx2* 1B2 over-expression (OE) construct; a −184 bp fragment including the duck “CA” SNPs; or a −184 bp fragment including the duck “CA” SNPs plus a 1B2 OE construct. Chick cells were also transfected with the duck (yellow) *Mmp13* promoter containing either a −181 bp control fragment including the duck “CA” SNPs; a −181 bp fragment including the duck “CA” SNPs plus a 1B2 OE construct; a −181 bp fragment including the chick/quail “AG” SNPs; or a −181 bp fragment including the chick/quail “AG” SNPs plus a 1B2 OE construct. In chick cells, 1B2 OE decreases activity of the chick and quail *Mmp13* promoters by 1.9-fold. Switching duck “CA” SNPs into the chick and quail promoters decreases activity by 1.7-fold and by 3.3-fold with 1B2 OE. Switching chick/quail SNPs into the duck promoter leads to an increase in activity by 1.5-fold both with and without 1B2 OE. **B)** In duck cells, 1B2 OE decreases activity of the chick *Mmp13* promoter by 1.7-fold and quail promoter by 1.5-fold. Switching the duck SNPs into the chick and quail promoters decreases activity by 3.8-fold even with 1B2 OE. **C)** In chick cells, 2B2 OE decreases activity of the chick *Mmp13* promoter by 2-fold and quail by 1.5-fold. Switching the duck SNPs into the chick promoter decreases activity by 2.5-fold and by 4.9-fold in the quail promoter. Switching chick/quail SNPs into the duck promoter leads to an increase in activity by 1.4-fold but when combined with 2B2 OE, duck promoter activity decreases. **D)** In duck cells, 2B2 OE decreases activity of the chick *Mmp13* promoter by 2.0-fold. Switching duck SNPs into the chick promoter decreases activity by 2.5-fold and by 11.2-fold in the quail promoter and also with 2B2 OE. Switching chick/quail SNPs into the duck promoter increases activity by 1.5-fold. Compared to the endogenous promoter within each species, * denotes significance at p ≤ 0.05, ** denotes significance at p ≤ 0.01, and *** denotes significance at p ≤ 0.001. # denotes significance at p ≤ 0.05 compared to control SNP switch. n = 18 to 36 for each treatment group.

**SUPPLEMENTAL TABLE 1.**
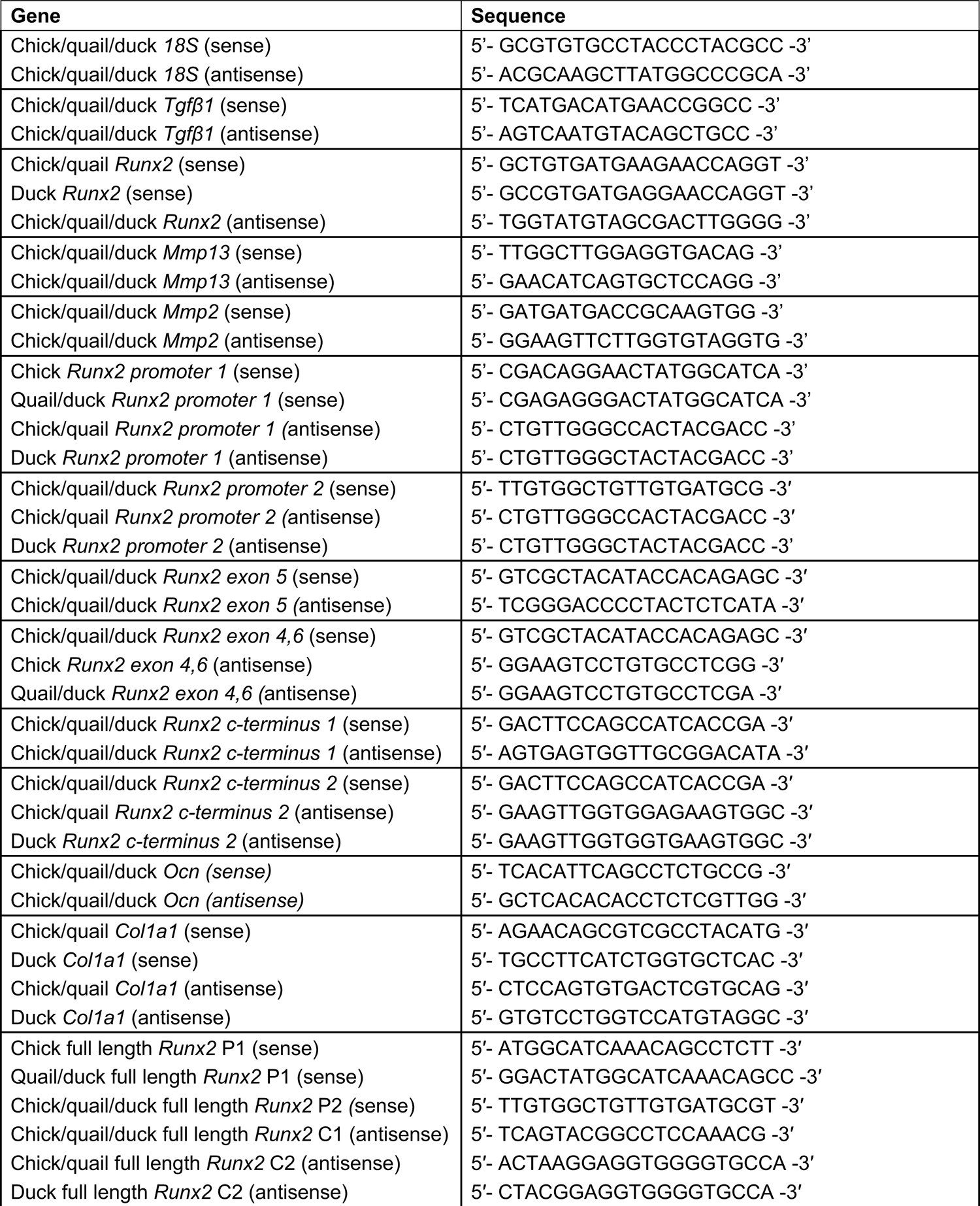
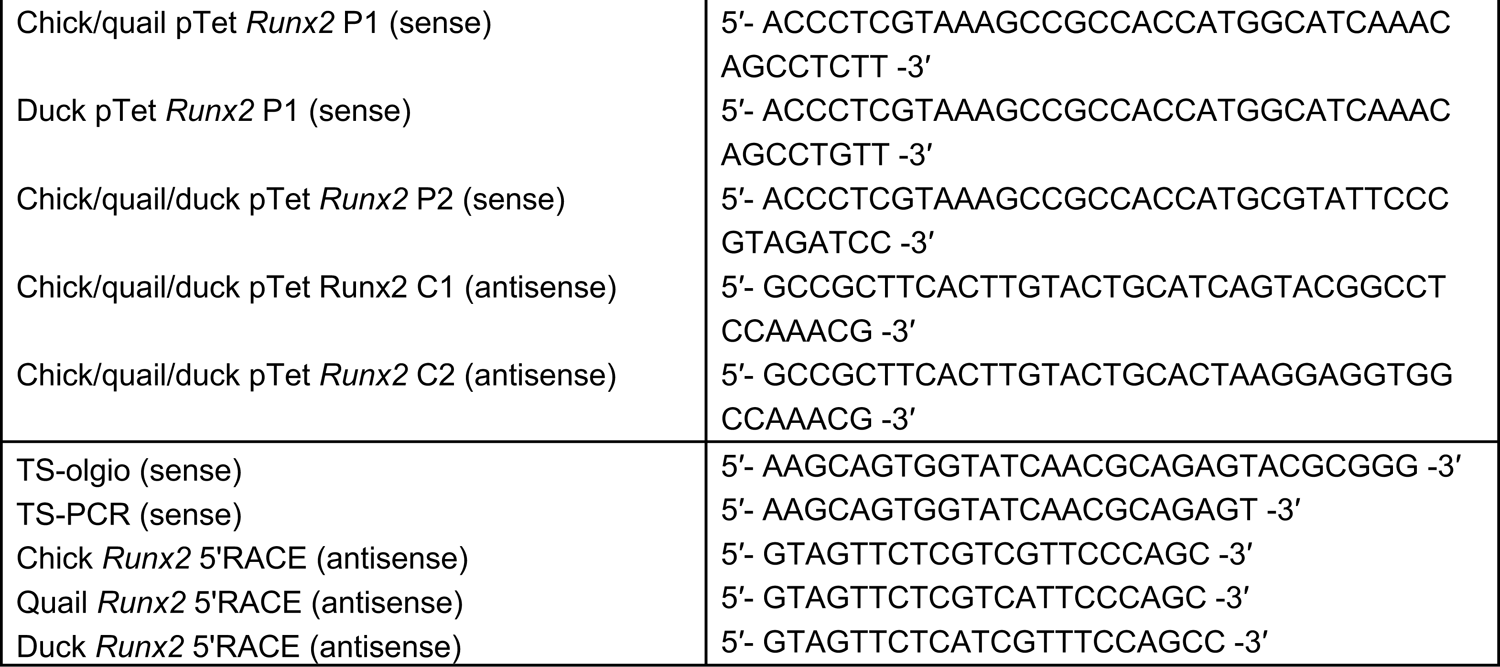
List of Primers

**SUPPLEMENTAL TABLE 2.**
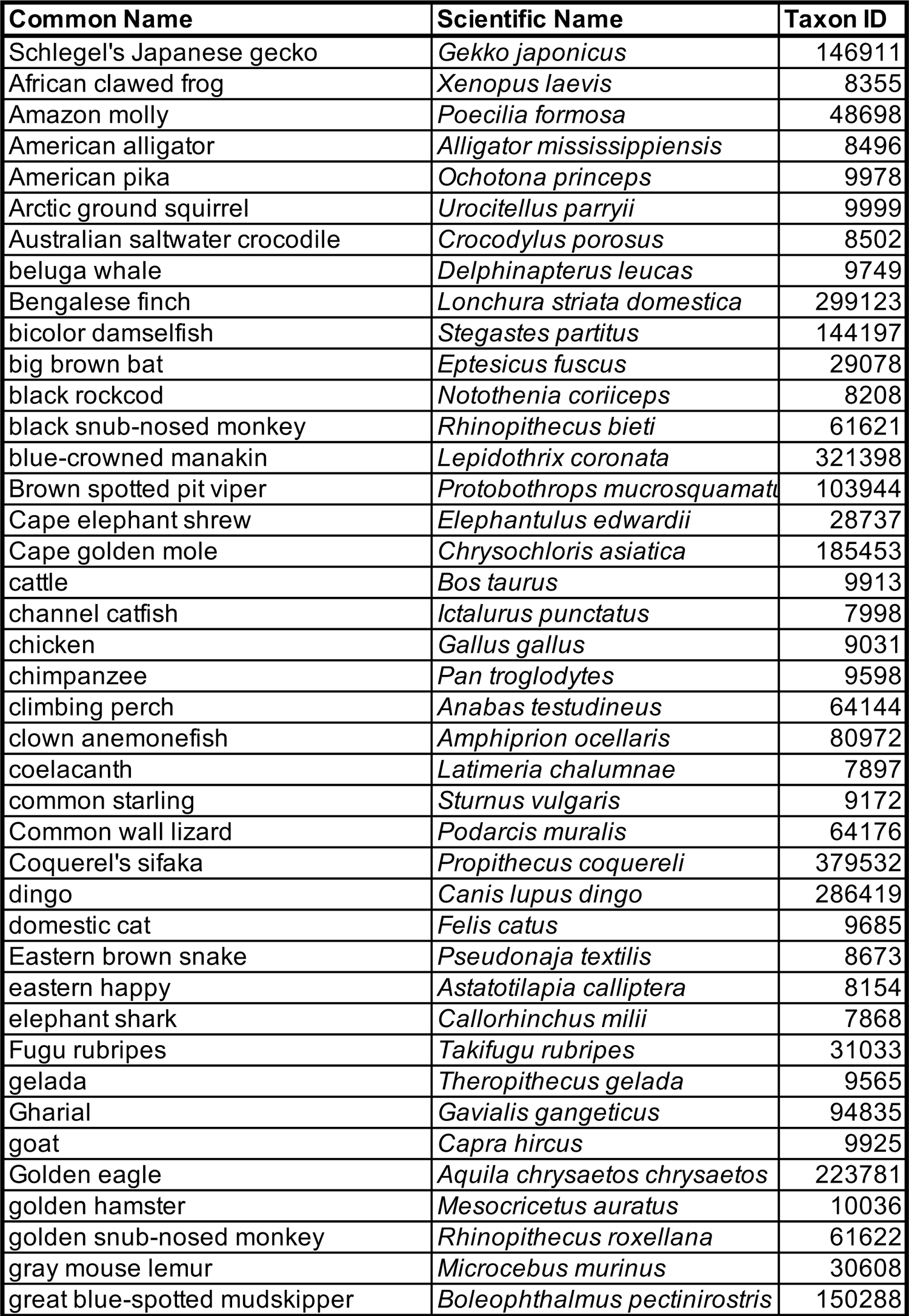
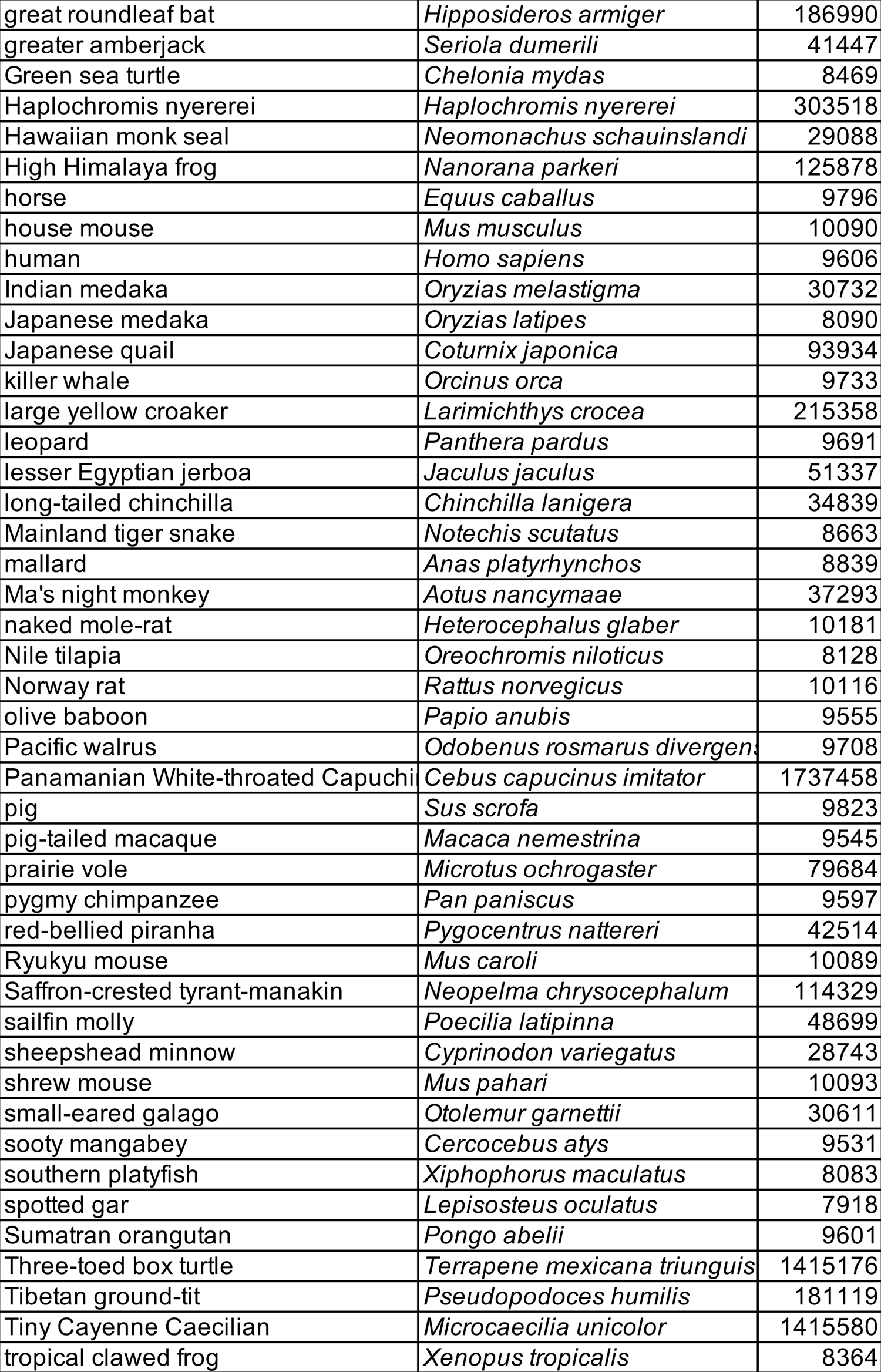
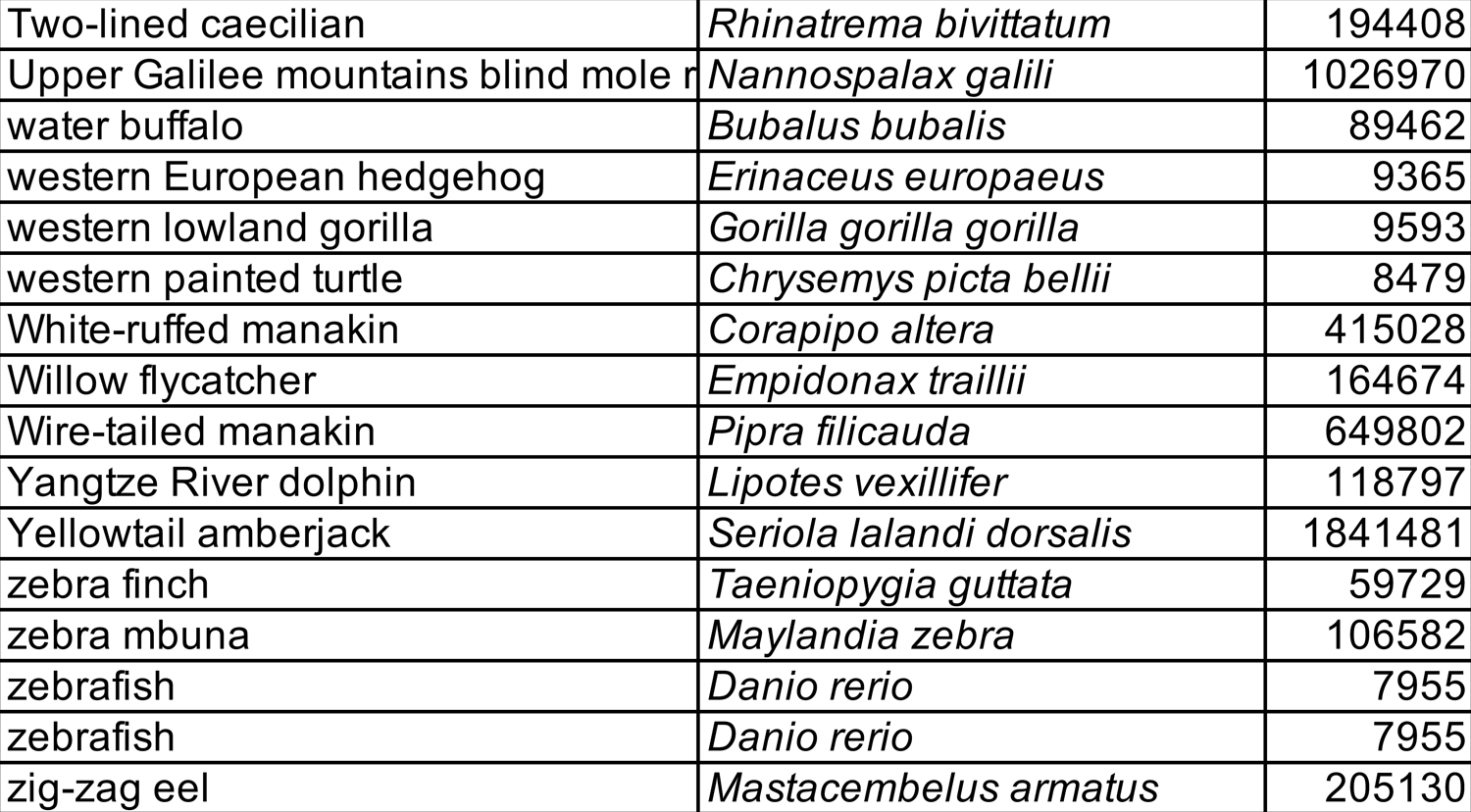
List of species from NCBI for which full-length RUNX2 was analyzed

**SUPPLEMENTAL TABLE 3.**
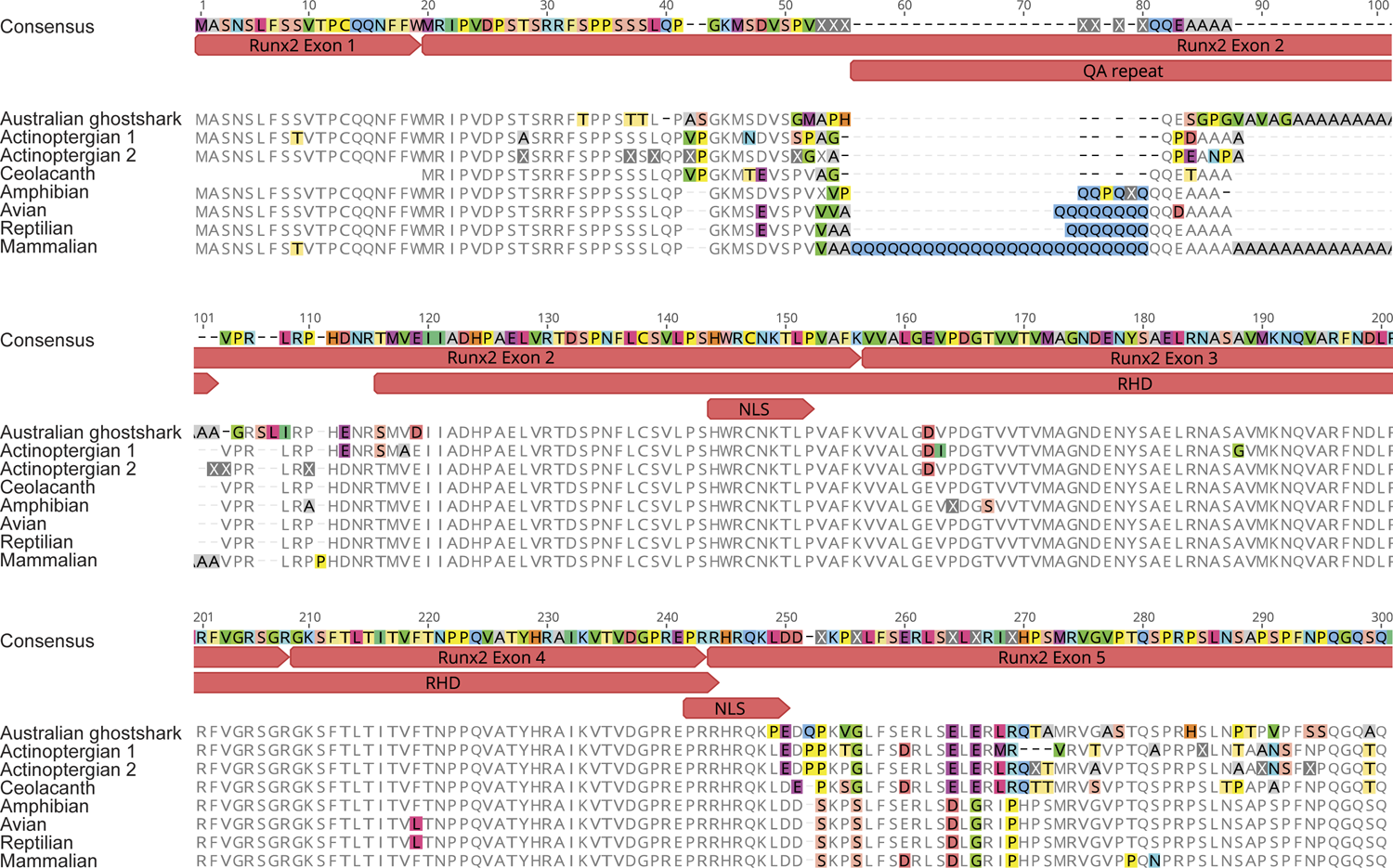
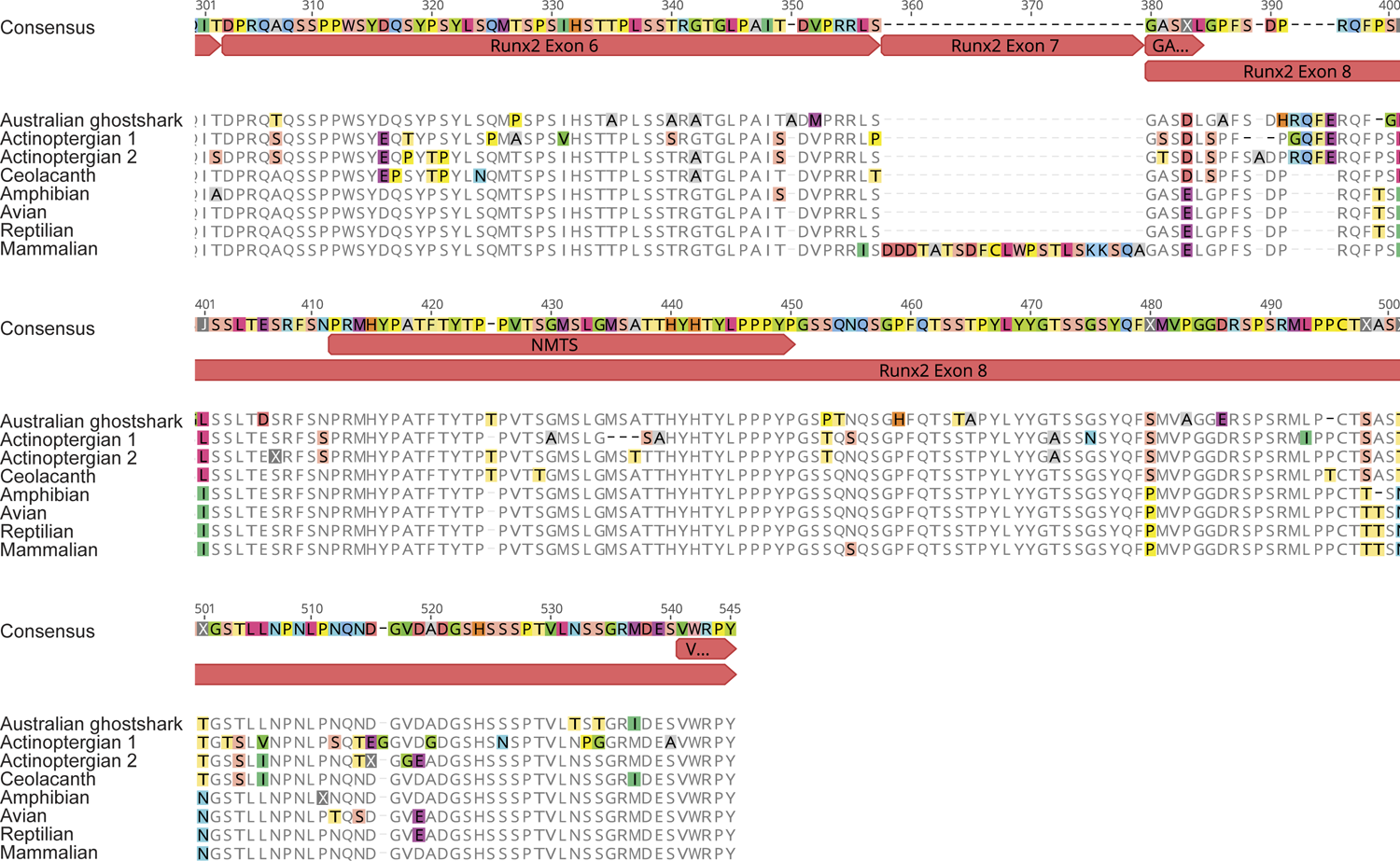
Consensus sequences from NCBI for full-length RUNX2 across vertebrates

**SUPPLEMENTAL TABLE 4.**
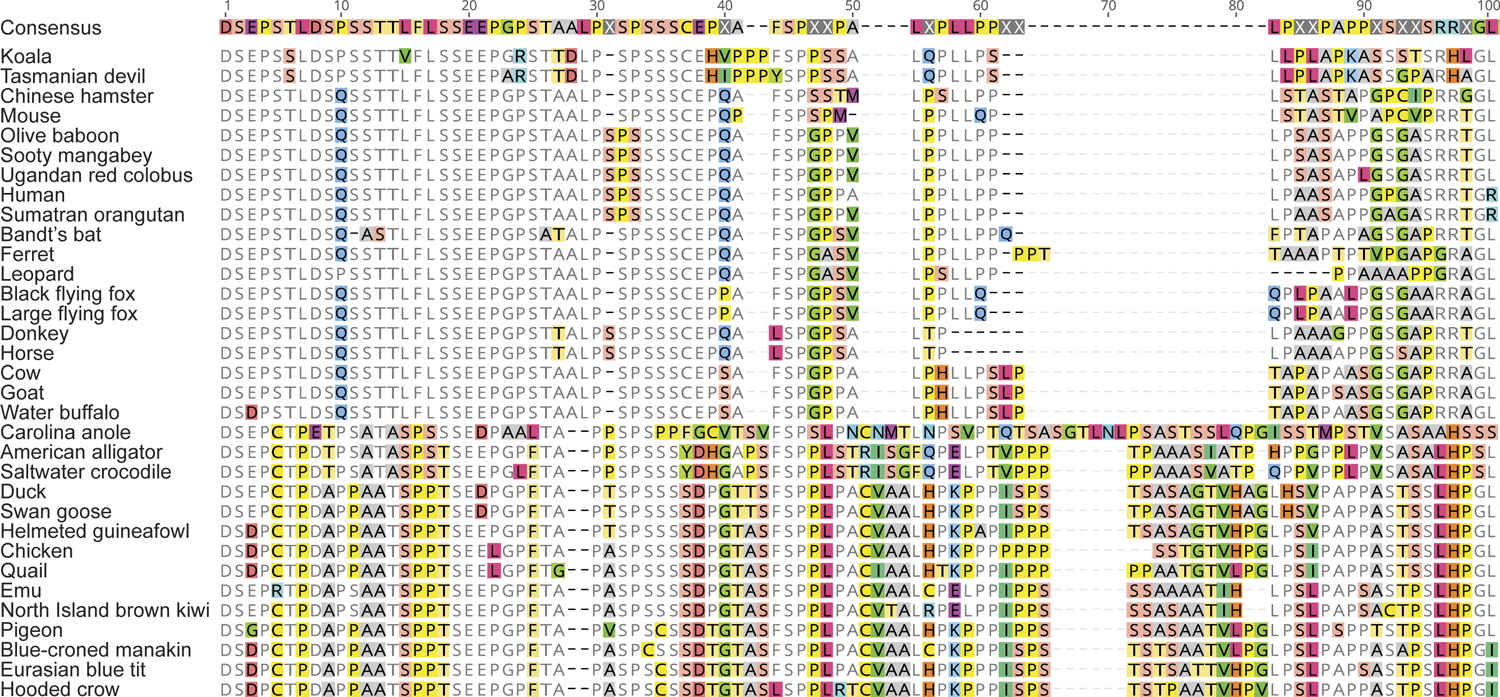
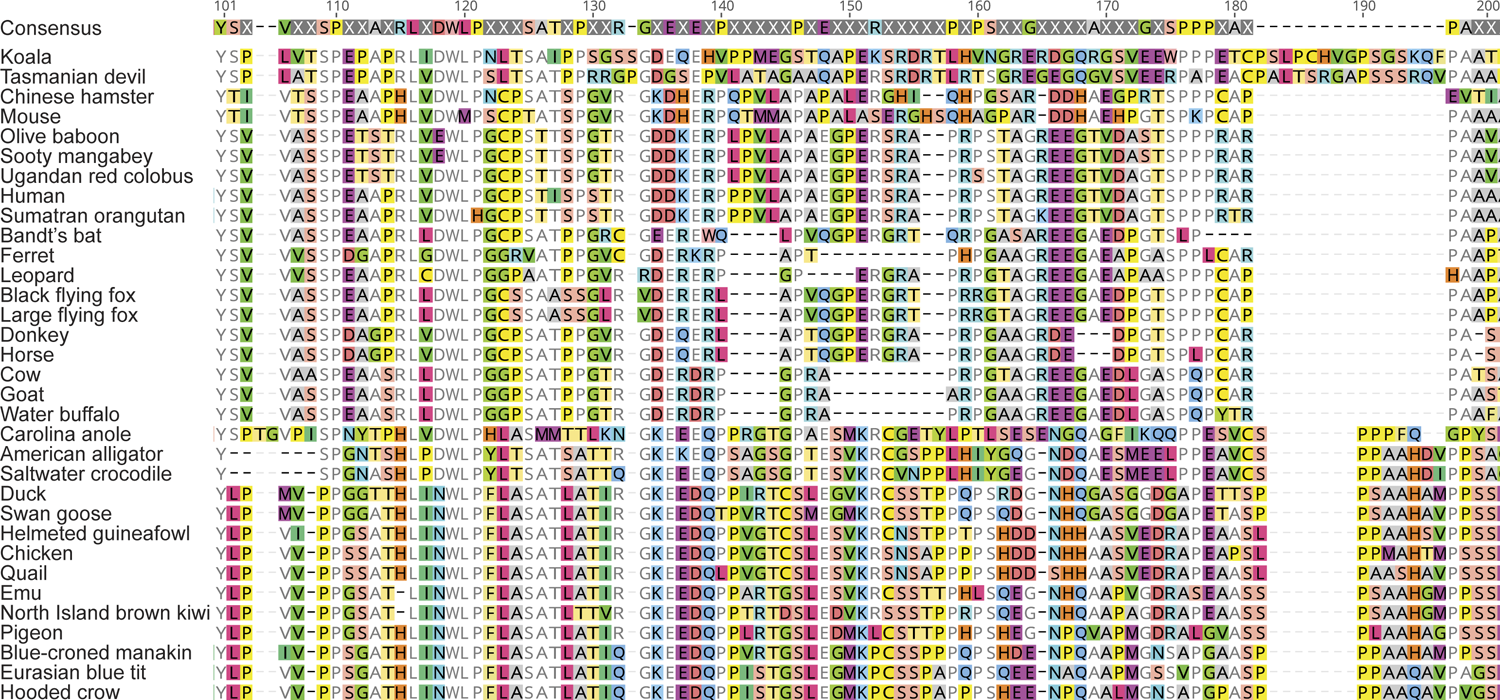
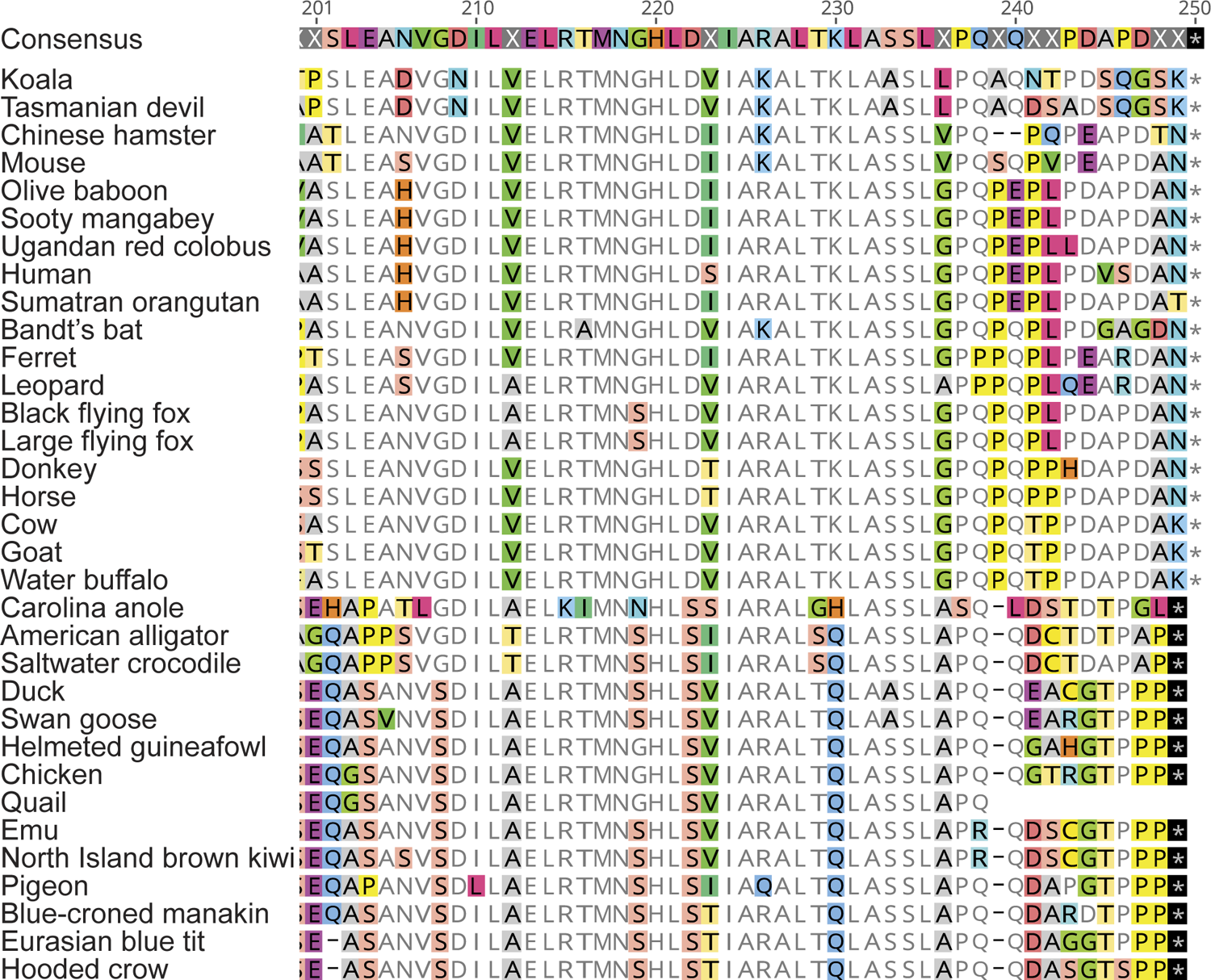
Consensus sequences from NCBI for C-Terminus 2 of RUNX2 in amniotes

